# Galbut virus infection minimally influences *Drosophila melanogaster* fitness traits in a strain and sex-dependent manner

**DOI:** 10.1101/2021.05.18.444759

**Authors:** Shaun T. Cross, Ali L. Brehm, Tillie J. Dunham, Case P. Rodgers, Grace I. Borlee, Mark D. Stenglein

## Abstract

Galbut virus (family *Partitiviridae*) infects *Drosophila melanogaster* and can be transmitted vertically from infected mothers or infected fathers with near perfect efficiency. This form of super-Mendelian inheritance should drive infection to 100% prevalence, and indeed galbut virus is ubiquitous in wild *D. melanogaster* populations. But on average only about 60% of individual flies are infected. One possible explanation for this apparent paradox is that a subset of flies are resistant to infection. Although galbut virus infected flies seem healthy, infection may be sufficiently costly to drive selection for resistant hosts, thereby decreasing overall prevalence. To test this hypothesis, we quantified a variety of fitness-related traits in galbut virus infected flies from two lines from the *Drosophila* Genetic Reference Panel (DGRP). Galbut virus infected flies had slightly decreased average lifespan and total offspring production, but these decreases were mostly not statistically significant. Galbut virus DGRP-517 flies pupated and eclosed slightly faster than their uninfected counterparts. Some galbut virus infected flies exhibited altered sensitivity to viral, bacterial, and fungal pathogens. Galbut virus infection produced minimal changes to host mRNA levels as measured by RNA sequencing, consistent with minimal phenotypic changes. The microbiome composition of flies was not measurably perturbed by galbut virus infection. Differences in phenotype attributable to galbut virus infection varied as a function of fly sex and DGRP strain and were generally dwarfed by larger differences attributable to strain and sex. Thus, galbut virus infection does produce measurable phenotypic changes, with changes being minor, offsetting, and possibly net negative.

**Importance:** Virology has largely focused on viruses that cause unmistakable phenotypic changes. But metagenomic surveys are revealing that persistent virus infections are extremely common, even in apparently healthy organisms. The extent to which these persistent viruses impact host fitness and evolution remains largely unclear. Here we study fitness impacts of a partitivirus named galbut virus that is ubiquitous in wild *D. melanogaster* populations. Despite efficient biparental vertical transmission, galbut virus is present in only just over half of wild flies. We quantified various fitness-related traits in galbut virus infected and uninfected fly lines and found that infection produced small but measurable changes in host phenotype that in aggregate may reduce fly fitness. Further studies that take advantage of this virus that naturally infects a premier model organism that is easy to study in the wild will shed further light on the persistent virus-host dynamics that may actually represent most viral infections.

## Introduction

Galbut virus is a remarkably successful persistent virus of *Drosophila melanogaster* [1–4]. Infected flies have been found on five continents and every wild population that has been tested includes some infected individuals. This degree of success is attributable to efficient biparental vertical transmission: infected mothers or infected fathers can transmit galbut virus to 100% of offspring, providing a means for infection to increase in frequency generation over generation [4,5].

Galbut virus belongs to a group of viruses, the partitiviruses (*Partitiviridae*), generally known for mild persistent infections [6–9]. Plant-infecting partitiviruses were originally called cryptic viruses (former genus *Cryptovirus*) because of their inapparent phenotypic effects [9,10]. Galbut virus infection is similarly cryptic: despite a century of *Drosophila* research and despite its ubiquity, galbut virus was only recently discovered by shotgun metagenomics [1]. Galbut virus infected flies do not exhibit obvious phenotypic differences from their uninfected counterparts, and in a population of wild-caught flies that we have maintained for three years, galbut virus has risen to and remained at 100% prevalence [4].

Nevertheless, there are indications that galbut virus may be in conflict with its host. Although infection is ubiquitous, the fraction of infected flies ranges from 13-100% in different populations, and on average only ~60% of flies are infected [1,3,4]. We also found that some *Drosophila* Genetic Reference Panel (DGRP) lines were relatively refractory to infection and multigenerational vertical transmission [4,11]. Galbut virus sequences exhibited high ratios of non-synonymous to synonymous variation (high dN/dS ratios), which could be consistent with selection driven by ongoing host-virus conflict [1]. Verdadero virus, a partitivirus that infects *Aedes aegypti*, exhibited similarly efficient biparental vertical transmission in colonized mosquitoes, but also is not at 100% prevalence in wild populations [12–14].

We hypothesized that galbut virus might exact a fitness cost that is small but sufficient to drive selection for resistant individuals. This could limit the overall success of galbut virus – and similar persistent viruses – in host populations. To test this hypothesis, we quantified a number of fitness-linked phenotypes in galbut virus infected flies. Host genotype and sex are variables that can substantially influence the outcome of infection [15–20], so we evaluated the phenotype of galbut virus infected males and females from two different DGRP strains.

Relatively little is known about arthropod-infecting partitiviruses, which have now been identified in association with a broad range of hosts including disease vectors [21–27]. Metagenomic surveys of apparently healthy free-living organisms have in general produced a flood of new virus sequences [12,28–30]. But beyond sequence description and phylogenetic placement, little is known about the biological impact of all of these newly recognized viruses. Follow-up virological studies that build upon this trove of sequence data are needed [4,27,31,32]. The ability to study a highly successful natural virus of a premier model organism represents a great opportunity to shed light on insect-infecting partitiviruses and persistent viral infections more generally.

## Methods and Materials

### *Drosophila* rearing and maintenance

Flies were reared on the Bloomington *Drosophila* Stock Center (BDSC) standard cornmeal diet (https://bdsc.indiana.edu/information/recipes/bloomfood.html). Stocks were housed at 25°C and changed every 14 days. All experiments were performed with *Drosophila* Genetic Reference Panel (DGRP) stocks 399 and 517, acquired from the BDSC [11,33]. Generation of galbut virus infected lineages by microinjection was described previously [4].

Experimental groups consisted of galbut virus infected or uninfected DGRP 399 or DGRP 517 males or females (2 strains x 2 sexes x 2 galbut virus infection status = 8 groups). All flies were 3-5 day old virgins reared in a 12 hour light/dark cycle at 25°C, unless otherwise stated.

### Quantification of galbut virus RNA levels

Total RNA was extracted from 5 day old, virgin flies using a bead-based protocol as previously described [4]. cDNA was synthesized by adding 5.5 μl of RNA to 200 pmol of a random 15-mer oligonucleotide and incubated for 5 min at 65°C, then set on ice for 1 min. A reverse transcription (RT) mixture containing the following was added (10 μL reaction volume): 1x SuperScript III (SSIII) FS reaction buffer (Invitrogen), 5 mM dithiothreitol (Invitrogen), 1 mM each deoxynucleotide triphosphates (dNTPs) (NEB), and 100 U SSIII reverse transcriptase enzyme (Invitrogen), then incubated at 42°C for 30 min, 50°C for 30 min, then at 70°C for 15 min. 90 μL of nuclease-free H2O was added to dilute the cDNA to a final volume of 100 μL.

### Quantification of major microbiome constituent DNA levels

Total DNA was extracted from 4-5 day old virgin flies. 10 flies per pool (total of 3 pools per group) were surface sterilized by vortexing in 70% ethanol for 2 minutes, followed by 2 rinses with autoclaved ddH2O and vortexing for 1 minute. Flies were then stored at −80°C until DNA was extracted. DNA was extracted using the DNeasy Tissue and Blood extraction kit (Qiagen) following the manufacturer’s protocol for insect tissues with three modifications. First, samples were added to 180 μL ATL buffer (provided in kit) along with a single BB bead and homogenized using a Qiagen TissueLyzer for 3 minutes at 30Hz rather than homogenizing by hand. Second, samples were incubated in proteinase K for a duration of 4 hours. Last, following incubation with proteinase K, samples were treated with 20 μL of RNase A (2 mg/mL; Sigma Aldrich) for 30 min at 37°C. After RNase treatment, samples were processed as stated in the protocol.

Following cDNA synthesis or DNA extraction, qPCR reactions were set-up using Luna qPCR Master Mix (NEB) following the manufacturer’s protocol. The qPCR reaction was performed on LightCycler 480 (Roche) under the following protocol: 95°C for 3 min, 40 cycles of 95°C for 10s, then 60°C for 45s, and then followed by a melting curve analysis. Microbiome analysis primer sequences were predominately from Early et al. [34]. Primer sequences can be found in **S1 Table**.

**Lifespan and fecundity assays** were performed similar to as previously described [35]. Flies were reared in 5 replicate groups of 10 adults (5 female, 5 male). Flies were checked daily for survival of adults, and living adults were moved to fresh media every 3 days. Longevity of adults was compared using the R survival package [36]. After adults were moved, original vials containing laid eggs were kept for 14 days, after which offspring were counted and sexed.

Total egg production was measured by housing 10 male and 10 female flies in bottles with an apple agar plate coated with yeast paste (1:1 yeast and water) to promote egg laying. Egg plates were replaced every 24 hours, and the used plates containing eggs were frozen at −20°C until the eggs were counted. Plates were collected for a total of 3 days, and were performed in 3 biological replicates and 2 technical replicates. Images of egg plates were captured and eggs were counted manually using the ImageJ cell counter program [37]. All fecundity measurements were analyzed with R scripts that can be found at:https://github.com/scross92/galbut_fitness_analysis.

**Developmental speed assays** were performed as previously described [38]. Eggs were collected using standard apple agar plates without preservative, with a mixture of 1:1 yeast and water applied. Every hour for 6-8 hours, agar plates were discarded and replaced to encourage egg synchronization. Agar plates were replaced a final time and incubated for several hours. The plates were removed and eggs were collected using an autoclaved brush. Twenty eggs were collected and moved to non-nutritive agar plates containing 5% sucrose/2% agar with no antimicrobials added (no tegosept). An agar plate was placed inside a larger petri dish with a damp paper towel on the bottom and moved to a 25°C incubator with a 12 hour light/dark cycle. Every 2 days, yeast paste was added as a nutrition source for developing flies. Yeast were killed prior to use in the paste by microwaving for 45 seconds on high to prevent overgrowth. Plates were checked daily for pupae to determine speed of pupation. Once pupation began, plates were checked approximately every 5 hours (morning, midday, evening). Continual monitoring occurred from pupation to emergence of adults in the same ~5 hour increments hours for measuring total time it took for flies to reach the adult stage. This was performed in 6 replicates per group (strain and galbut virus infection status).

### *Pseudomonas aeruginosa* oral challenge

Flies were challenged orally with *Pseudomonas aeruginosa* as adapted from Lutter et al [39]. An overnight culture of *P. aeruginosa* (PAO1) was grown in a 200 mL culture Brain Heart infusion (BHI) broth incubated at 220 rpm at 37°C. The following day, the culture was centrifuged at 4200 g for 5 minutes until a loose pellet was formed. Excess supernatant was decanted and culture was resuspended to an OD_600mm_ of ~7 using a sterile 5% sucrose solution. Autoclaved filter disks were inoculated with 290 μL of the *P. aeruginosa* solution. Disks were placed on 5% sucrose agar vials. Control disks were inoculated with the 5% sucrose solution. Twelve flies that had been starved for 5 hours were placed in the bacteria-containing vials for each replicate. Flies that died by the end of the first day were censored from further analysis, since their deaths were likely due to starvation stress. Survival of flies was monitored daily for 12 days. Statistical analysis was performed using the R survival package [36]. A total of 3 technical replicates were performed.

### Intrathoracic microbial pathogen challenges

The following pathogen challenges were performed through intrathoracic microinjection. All experimental injections were performed in 3 biological replicates (12 flies per replicate) per technical replicate, and a total of two technical replicates were performed for each pathogen. An exception is the *Staphylococcus aureus* challenge which was performed in 3 technical replicates. Control injections with 1x phosphate buffered saline (PBS) were performed in parallel. Flies were checked at 10-12 hours post-injection, and any flies that were dead at this point were assumed to have died from injection. Additionally, any flies that died from non-natural causes (for example, after getting stuck in the media) were also censored from analysis. Injected volumes, inoculum dose, and subsequent intervals for checking fly survival are stated below for the respective pathogen.

#### Pseudomonas aeruginosa

Flies were microinjected with *P. aeruginosa* (strain PAO1). A culture was started by inoculating 150mL of BHI broth and incubated at 220 rpm overnight at 37°C. The following day, the culture was centrifuged at 4200 g for 5 minutes until a loose pellet was formed. Excess supernatant was decanted and the culture was resuspended to an OD_600mm_ of 0.03 using 1x PBS. Flies were injected with 9.2 nL of this diluted *P. aeruginosa* culture, which corresponds to ~100 CFUs [40]. Flies were incubated overnight and checked at 24 hours post injection, 28 hours post injection, and every 2 hours from 28 to 42 hours post injection. After 42 hours post injection, flies were checked at one final time point of 52 hours post injection, at which any living flies were censored from downstream statistical analyses.

#### Staphylococcus aureus

Flies were microinjected with *S. aureus* (strain XEN36, Perkin Elmer). A culture was obtained by inoculating 150mL BHI broth and stirred at 220 rpm overnight at 37°C. The following day, the culture was centrifuged at 4200 g for 5 minutes until a loose pellet was formed. Excess supernatant was decanted and the culture was resuspended to an OD_600mm_ of 0.1 using 1x PBS. Flies were injected with 23 nL of this diluted *S. aureus* culture, which corresponds to ~100 CFUs [41]. Flies were checked daily until 8 days post injection, at which point any living flies were censored from downstream statistical analyses.

#### Drosophila C virus

*Drosophila* C virus (DCV) stocks were provided by the Andino lab at the University of California San Francisco. DCV stocks were amplified and titrated on *Drosophila* S2 cells. DCV infections of flies were performed as previously described [42]. Flies were microinjected with DCV at a titre of 100 50% Tissue Culture Infective Dose units (TCID50) in a total volume of 50 nL. Flies were checked daily until 14 days post injection, at which point any living flies were censored from downstream analyses.

#### Candida albicans

*Candida albicans* challenge was performed as previously described [43]. *C. albicans* (strain SC5314) was obtained from ATCC. A yeast extract peptone dextrose (YPD) agar plate was streaked from the frozen glycerol stock and incubated at 30°C for 18 hours. 150mL of YPD broth was inoculated with a single colony from the YPD plate and incubated at 220 rpm overnight at 30°C until the culture was at an OD_600mm_ of ~1. The culture was centrifuged at 4200 g for 5 minutes until a loose pellet was formed, which was resuspended using 1x PBS. Yeast cells were counted with a cytometer and diluted to 10^7^ cells/mL. Flies were microinjected with 50 nL (~500 cells) of this dilution. Flies were incubated at 30°C and were checked daily until 6 days post injection, at which point any living flies were censored from analyses.

### RNAseq library preparations

Pools of 10, 5-day old, virgin flies were collected, flash frozen in liquid nitrogen and stored at −80°C. Polyadenylated RNAs were enriched using the NEB Magnetic mRNA Isolation Kit according to the manufacturer’s protocol. Sequencing libraries were created using Kapa HyperPrep RNA Library Prep Kit (Roche) according to manufacturer’s protocol. Final library molecules had an average size of 348 base pairs, and were sequenced on an Illumina NextSeq using the NextSeq 500/550 High Output Kit v2.5, generating 75 base single-end reads.

### Transcriptomic computational analysis

RNAseq datasets were first processed to remove low quality and adapter sequences using cutadapt tool [44] version 1.13 with the following settings: -a AGATCGGAAGAGC -A AGATCGGAAGAGC -g GCTCTTCCGATCT -G GCTCTTCCGATCT -a AGATGTGTATAAGAGACAG -A AGATGTGTATAAGAGACAG -g CTGTCTCTTATACACATCT - G CTGTCTCTTATACACATCT, -q 30,30, --minimum-length 40, and -u 1. Remaining reads were mapped to the *D. melanogaster* genome assembly BDGP6.28 from Ensembl using HISAT2 version 2.2.0 [45]. Read mapping was tabulated using featureCounts version 2.0.0 [46] to the BDGP6.28 gtf file with the following settings: -s 2 -t exon -g gene_id. The resulting read count table was used as input for differential gene expression analysis using DESeq2 version 1.26.0 [47] in R version 3.6.3 [48]. Differential gene expression analyses on the condition of galbut virus infection status and was performed for each group (strain and sex). Gene set enrichment analyses (GSEA) were performed using the clusterProfiler R package version 3.14.3 [49] using a pre-ranked gene list ordered by the log2 fold changes and the ‘gseGO’ function. Redundant GO terms were collapsed using the ‘simplify’ function by adjusted p values, with a cutoff value of 0.7, and the “Wang” measure.

### Data Availability

Computational scripts for analysis of experiments can be found at https://github.com/scross92/galbut_fitness_analysis. Raw sequencing data can be found on the NCBI Sequence Read Archive (SRA) under Bioproject PRJNA683038.

## Results

### Confirmation of galbut virus infection status and galbut virus RNA levels in individual flies

We first verified that our stocks of galbut virus-infected flies established previously were still persistently infected [4]. We quantified galbut virus RNA levels using qRT-PCR in 20, 3-5 day old flies from each line (10 male and 10 female), and normalized levels to those of ribosomal protein L32 mRNA (RpL32; **Fig 1**). Galbut virus RNA levels were higher than those of highly expressed RpL32 mRNA in all cases (**Fig. 1**). Median galbut virus RNA levels were 2.3-fold higher in DGRP 399 flies than in DGRP 517 flies (p=1.6×10^-2^), although some outlier DGRP 517 flies had higher levels (**Fig 1**). Galbut virus RNA levels were 2.1-fold higher in DGRP 399 males than in females (p=4.2×10^-5^) and 1.5-fold higher in DGRP 517 males than in females (p=1.3×10^-2^). So, these populations remained persistently infected at 100% prevalence, and galbut virus RNA levels varied as a function of DGRP strain and sex, with levels generally higher in males and in DGRP 399 flies.

**Fig 1.**
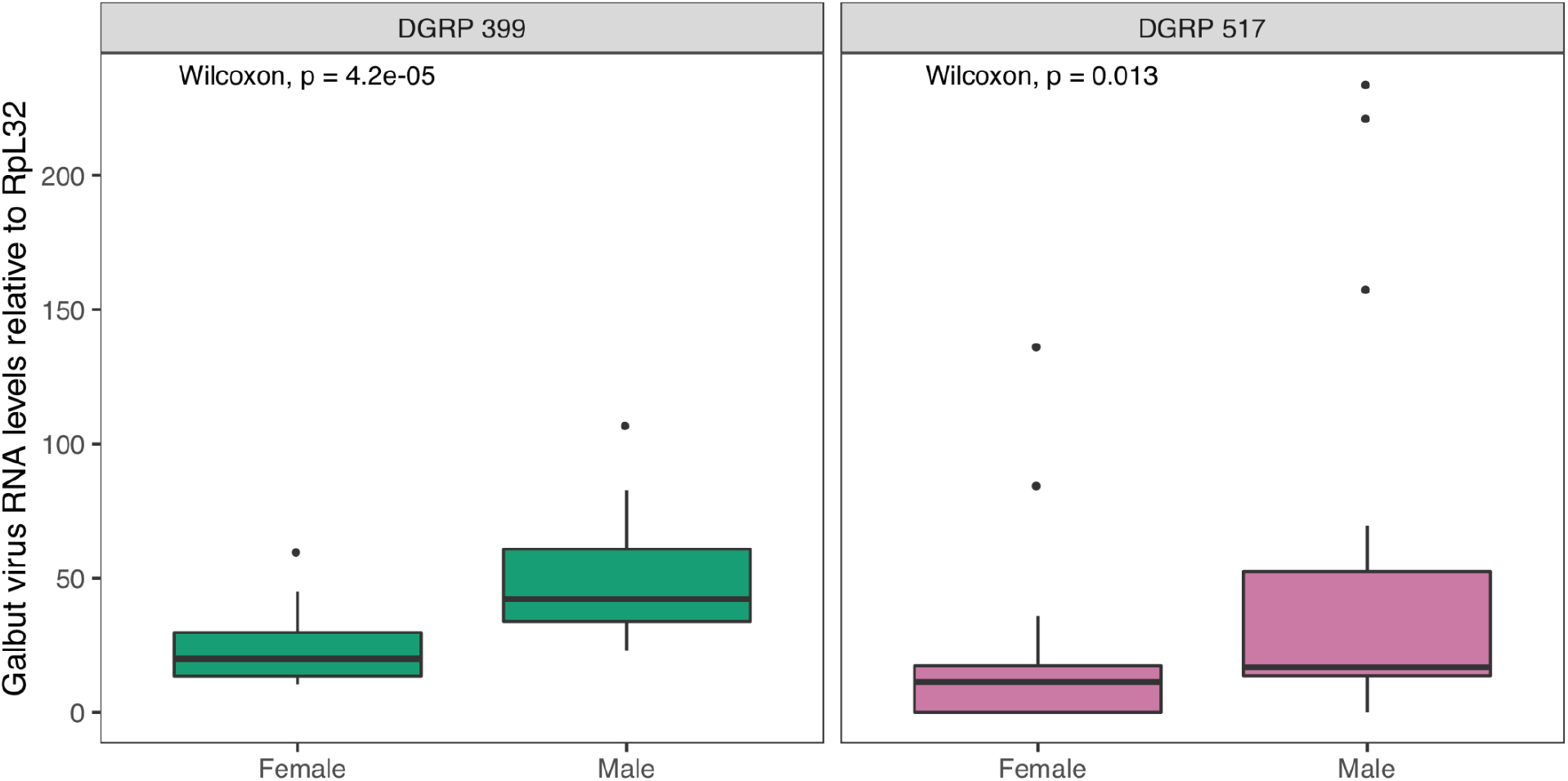
Relative galbut virus RNA levels in flies from two DGRP strains. Boxplots depicting galbut virus RNA levels relative to the housekeeping gene RpL32 (2^-ΔCt^ method) in DGRP 399 and DGRP 517 adult flies from strains used in this study (n = 10 per strain and sex).

### Galbut virus infection does not have significant impacts on predominant microbiome constituents

The microbiome composition of *D. melanogaster* can alter fitness [35]. It’s also possible that viral and bacterial constituents of the microbiota can interact [50]. Commensal bacteria can also vary by DGRP background when reared under the same conditions [34]. We therefore tested whether the microbiomes of these populations varied as a function of galbut virus infection status. Our goals were to assess whether microbiome differences could underlie differential phenotypes in flies with and without galbut virus, and whether galbut virus infection was altering microbiome composition.

Previous shotgun metagenomic RNA sequencing of DGRP 399 and 517 flies reared in our lab had revealed that the major constituents of these flies’ microbiomes included *Acetobacter persici*, *Lactobacillus brevis*, *L. planatarum*, *Corynebacterium* spp., and *Saccharomyces cerevisiae*. We quantified DNA levels of these microbes by qPCR using previously designed primers. DNA copy numbers were normalized to the single copy host gene *deformed* (*dfd*) as previously described [34]. The relative abundances of the different microbes were similar in DGRP 399 and 517 flies and in males and females (**Fig. 2**). Galbut virus infection did not produce any statistically significant differences in DNA levels of these taxa in any of the groups (**Fig 2**). This indicated that galbut virus infection did not measurably change microbiome composition and that any fitness effects of galbut virus infection were likely not mediated by changes in microbiome composition.

**Fig 2.**
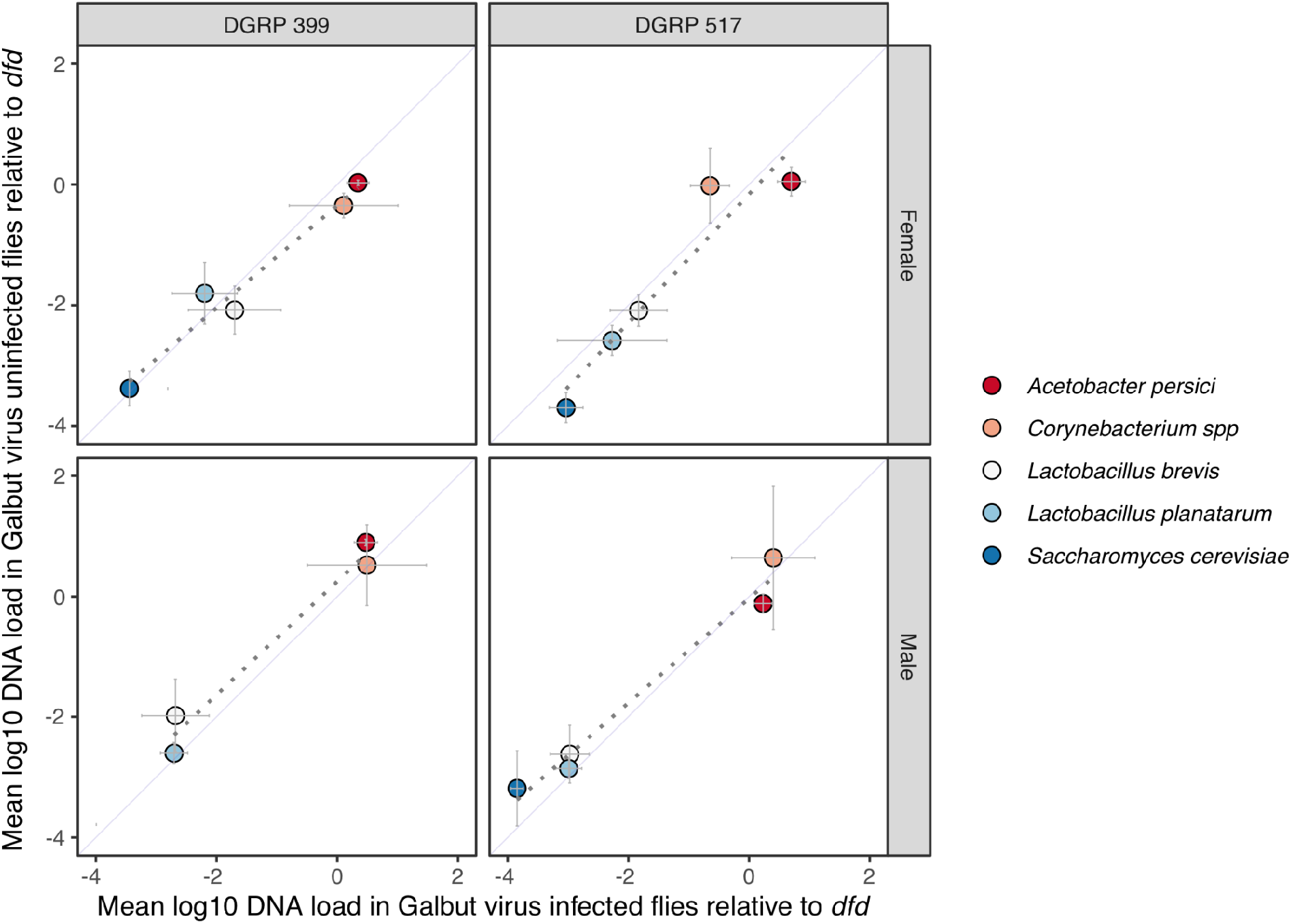
Galbut virus infection did not alter levels of major microbiome constituents in flies. Relative amounts of DNA from predominant microbiome constituents in galbut virus infected and uninfected flies were measured via qPCR from 3 replicate pools of 10 flies per pool per strain per sex. Mean DNA loads relative to single copy *dfd* gene are plotted and crossbars indicate standard deviations of replicates. Dotted lines depict linear regression fits and solid lines indicate the diagonal. No statistically significant differences between galbut virus infected and uninfected flies were identified using a Wilcoxon test.

### Galbut virus slightly reduces *Drosophila* lifespan and fecundity

We compared the lifespan, fecundity, and developmental speed of galbut virus infected and uninfected flies [35,51]. Vials of newly eclosed adults (n=5 replicate vials per experimental group) were housed together in groups of 10 flies (5 males, 5 females). Galbut virus infected flies from both strains exhibited a slightly shortened mean lifespan (6.6 days shorter in DGRP 399 and 3.1 days shorter in DGRP 517), though this was not statistically significant in either strain (**Fig 3A**). These decreases in lifespan attributable to galbut virus infection status were smaller than differences attributable to DGRP strain: DGRP 517 flies lived on average 14.9 fewer days than DGRP 399 flies (p=2.1×10^-4^, **Fig 3B**).

**Fig 3.**
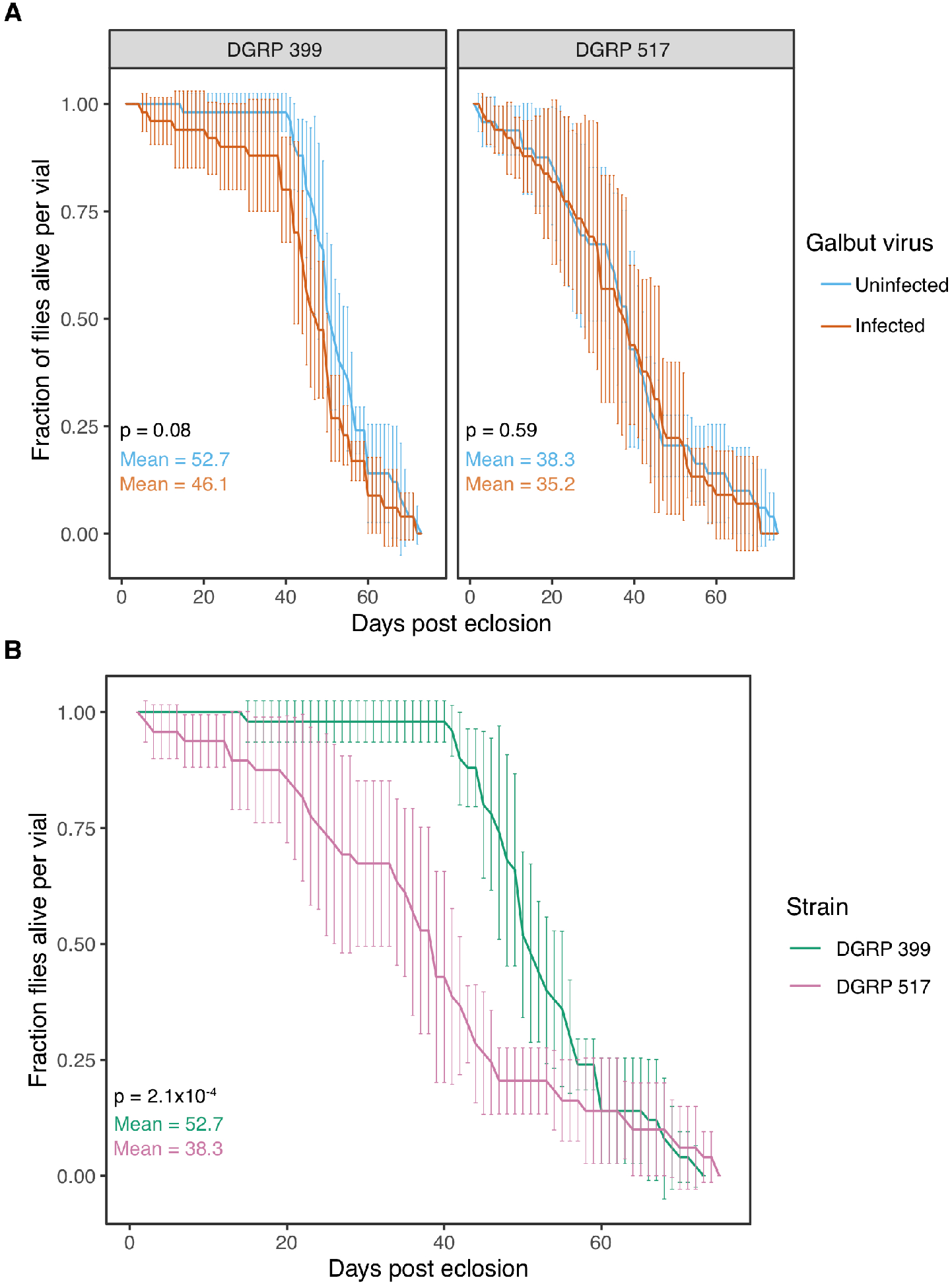
Galbut virus infected flies exhibit slightly reduced lifespan. (A) Survival of DGRP 399 and 517 flies with or without persistent galbut virus infection. The mean and standard deviation of biological replicates is plotted. (B) Data as in A, but plotted to facilitate comparison of DGRP strains.

We compared fecundity of infected and uninfected flies by counting total adult offspring in vials containing 5 male and 5 female flies. Galbut virus infected DGRP 399 flies produced fewer offspring than their uninfected counterparts, but this was not significantly different (t-test; female offspring p=0.77, male offspring p=0.83; **Fig 4A**). Galbut virus infected DGRP 517 flies also produced fewer offspring, but the decrease was only significant for male offspring numbers (t-test; female offspring p=0.16, male offspring p=0.027; **Fig 4A**). The total offspring counts were not normalized to surviving mothers, so the lower number of offspring at later timepoints likely reflect the slightly shortened lifespan of galbut virus infected flies (**Fig 3**). Galbut virus infection did not significantly change offspring sex ratios (t-test; DGRP 399: p=0.63, DGRP 517: 0.75, **Fig. S1**). As with average lifespan, differences in total offspring number between the different DGRP strains were much larger than differences attributable to galbut virus infection status: DGRP 399 females produced on average 2.7x more offspring than DGRP 517 females (p=2.8×10^-6^, **Fig 4B**).

**Fig 4.**
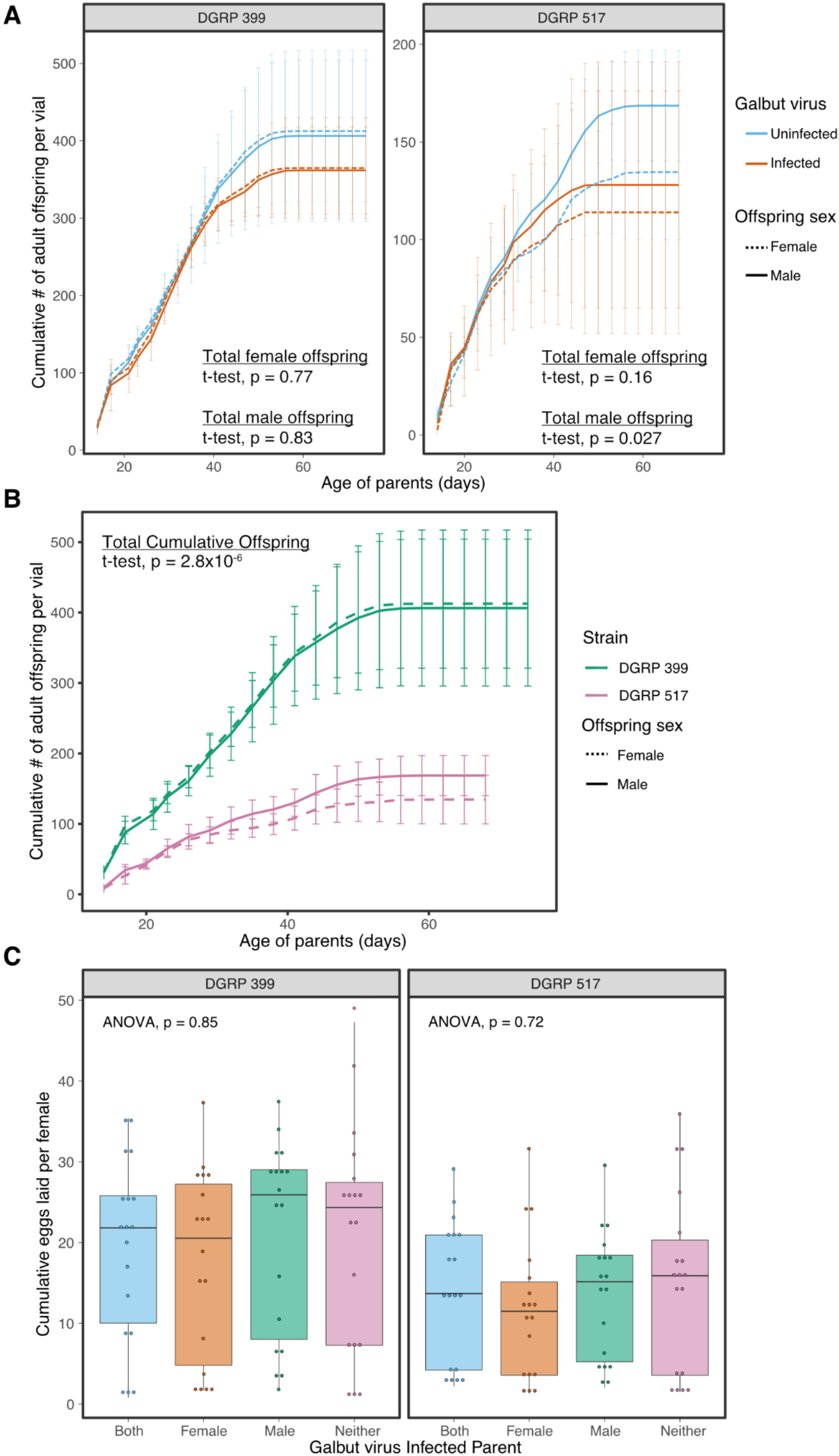
Galbut virus infected flies produce slightly fewer offspring. (A) Galbut virus infected and uninfected flies were housed in batches of 5 males and 5 females per vial and cumulative number of female and male offspring per vial were counted. The mean and standard deviation of biological replicates are plotted. (B) Data as in panel A, but plotted for comparison of DGRP strains (data from galbut virus uninfected flies shown). (C) 10 male and 10 female flies 3-5 days post eclosion were crossed with different combinations of galbut virus infected mothers or fathers. The cumulative number of eggs laid per female over three days is depicted for individual replicates as points and summarized with boxplots.

We recorded the cumulative number of eggs laid over three days when one or both parents were infected by galbut virus. There were no significant differences in the number of eggs laid when either or both parents were infected with galbut virus (ANOVA, DGRP 399: p=0.85, DGRP 517: p=0.72; **Fig 4C**).

We compared the developmental speed of galbut virus infected and uninfected flies by collecting eggs and monitoring the times from oviposition to pupation and oviposition to adulthood. DGRP 399 flies pupated in ~5 days and eclosed in ~9 days regardless of galbut virus infection status (**Fig 4A & 4C**). DGRP 517 flies infected with galbut virus pupated on average 7 hours faster than uninfected flies (Wilcoxon, p=2.2×10^-16^; **Fig 4A**). DGRP 517 infected females and males reached adulthood on average 10 and 12 hours faster than their uninfected counterparts (**Fig 4C**). As was the case with other phenotypes, development speed also varied as a function of DGRP background, with DGRP 399 flies pupating on average 7 hour faster than DGRP 517 flies (**Fig 4B**) and eclosing on average 13 hours faster (**Fig 4D**).

### Galbut virus alters the susceptibility of flies to viral, bacterial, and fungal pathogens

The microbiota present in a host can influence the outcome of subsequent infections [52–55]. Moth-infecting partitiviruses changed their host’s ability to withstand infection by a pathogenic nucleopolyhedrovirus [27]. We hypothesized that galbut virus infection might alter the ability of flies to resist or tolerate infection by pathogenic microbes, which could alter the survival and consequently the fitness of galbut virus infected flies. To test this hypothesis, we challenged galbut virus infected and uninfected flies with viral, bacterial, and fungal pathogens.

We first tested whether pre-existing galbut virus infection altered fly survival following infection by *Drosophila* C virus (DCV) [56]. Flies were challenged with 100 TCID_50_ units of DCV through intrathoracic microinjection and checked daily for survival. Overall, there was little difference in the survival of galbut virus infected and uninfected flies. DGRP 517 female galbut virus-infected flies survived slightly longer than their uninfected counterparts, and although this effect was statistically significant, it was small in magnitude (**Fig 6A**, p=0.028). These DGRP strains are both *Wolbachia* negative, so improved survival could not be attributed to the known protective effects of *Wolbachia* against DCV [57–60].

**Fig 5.**
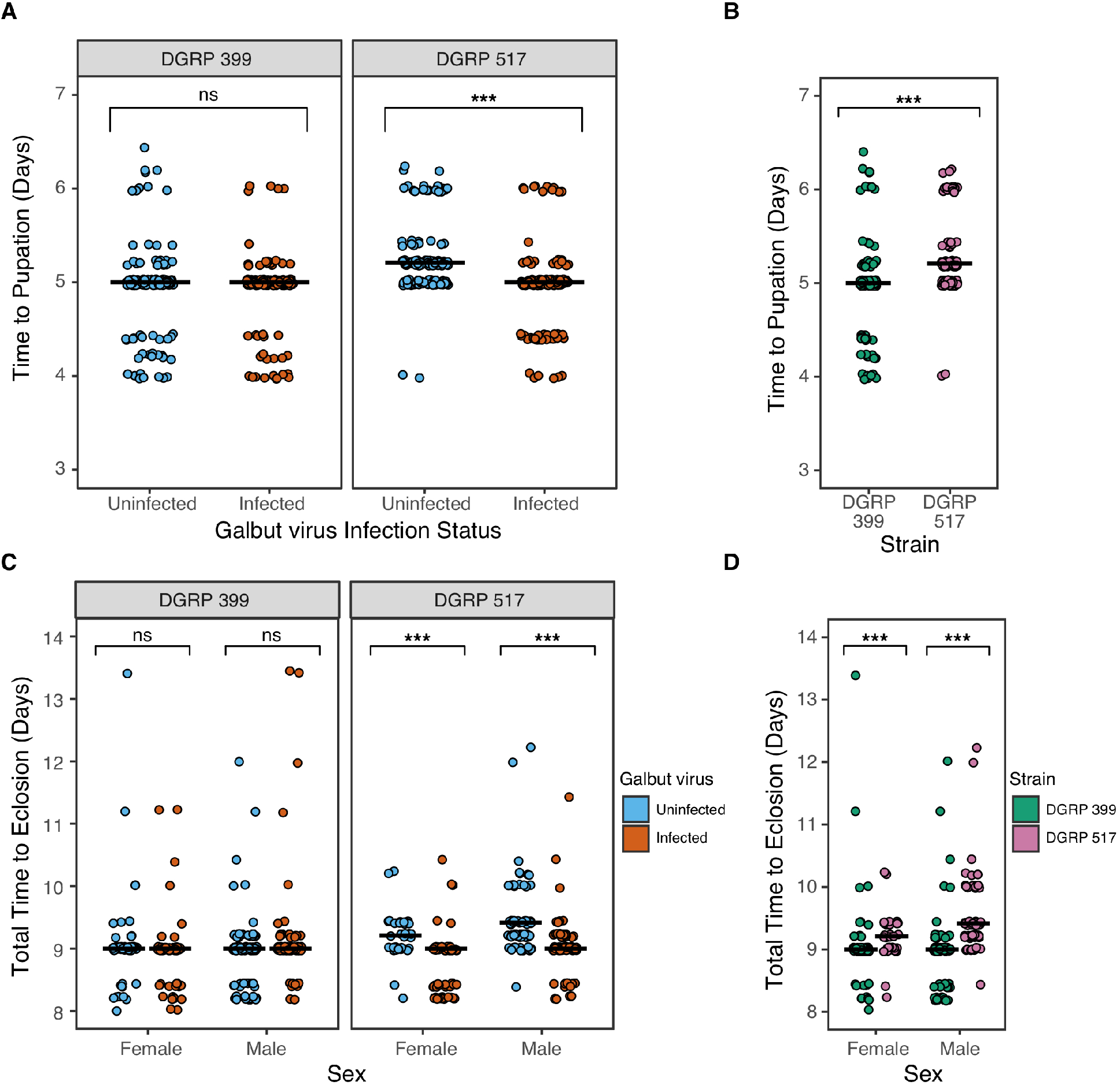
Galbut virus infected DGRP 517 flies develop slightly faster. (A) The time to pupation of individual DGRP 399 or DGRP 517 flies is plotted and the median time is indicated by a crossbar. (B) Data as in A, but plotted to enable comparison between DGRP strains. (C) The time between oviposition and eclosion for individual flies is indicated. (D) Data as in C, but plotted to enable comparison between DGRP strains. ns: not significant; **: p < 0.01, ***: p < 0.001

**Fig 6.**
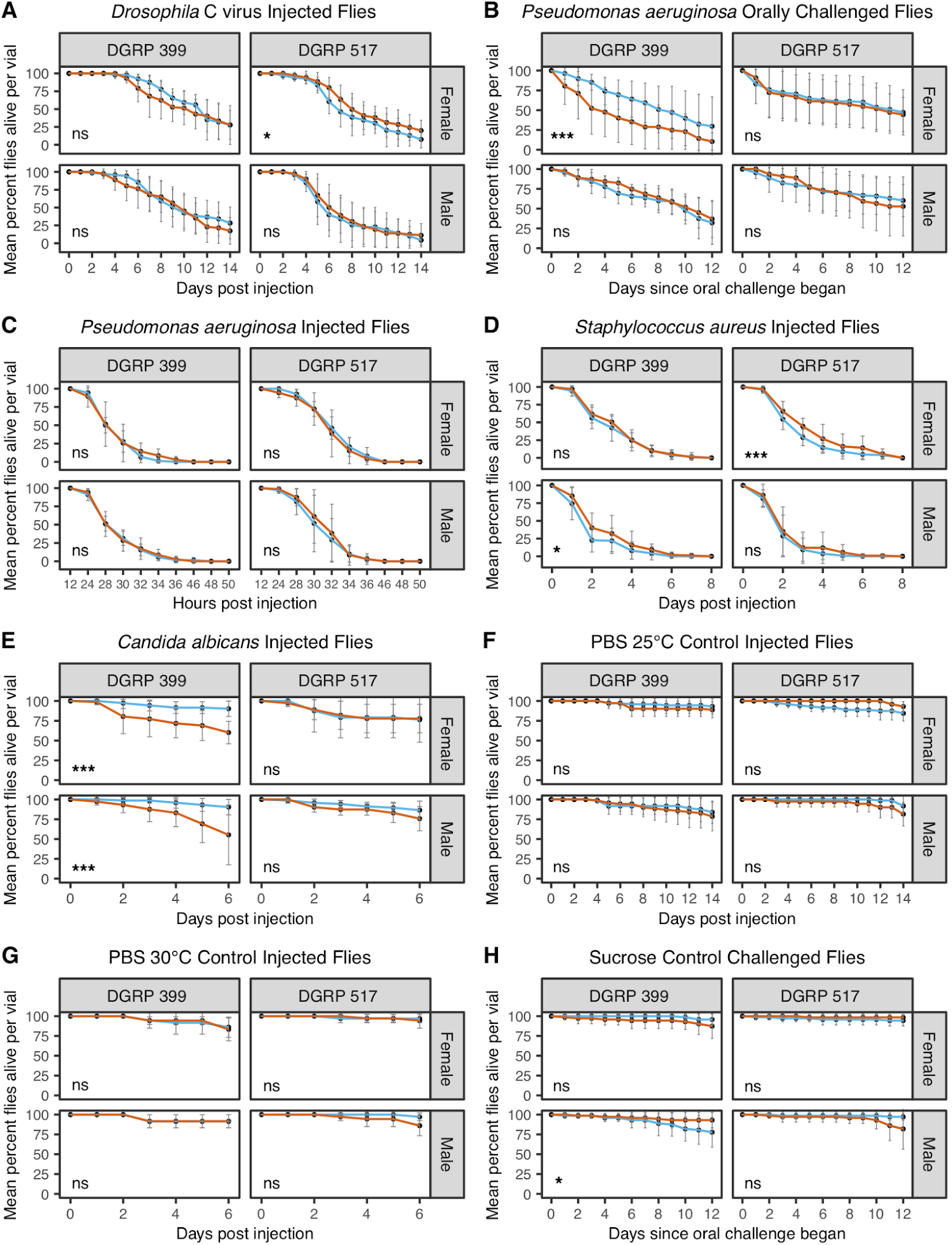
Galbut virus alters pathogen susceptibility of some flies. Survival of galbut virus infected and uninfected flies following (A) intrathoracic injection with 100 TCID_50_ units of *Drosophila* C virus (DCV), (B) ingestion of *Pseudomonas aeruginosa*, (C) injection of ~100 CFUs of *Pseudomonas aeruginosa*, (D) injection with ~100 CFUs of *Staphylococcus aureus*, (E) injection with ~500 *Candida albicans* cells. (F-H) Survival of flies following control inoculations. Flies were either microinjected with phosphate buffered saline (PBS) and stored at 25°C (F) or 30°C (G), or ingested sucrose (H). Galbut virus infected flies are depicted in orange and uninfected flies are in blue. ns: not significant; *: p < 0.05; **: p < 0.01, ***: p < 0.001.

We next challenged flies orally with *Pseudomonas aeruginosa*. Galbut virus infected DGRP 399 female flies were more susceptible to *P. aeruginosa* bacterial challenge (**Fig 6B**; p=4.5×10^-6^). Although ingestion is a more natural route of infection than microinjection, there is less experimental control over the ingested dose, which can decrease reproducibility [40]. We therefore also injected flies with ~100 CFUs of *P. aeruginosa*. Flies injected with *P. aeruginosa* died faster than those that ingested the pathogen, with most flies dead by 36 hours post injection (**Fig. 6C**). Galbut virus infected DGRP 399 females no longer died faster than their uninfected counterparts when microinjected with *P. aeruginosa* (**Fig 6C**; p=0.14). This suggests that interactions between galbut virus and *P. aeruginosa* may depend on the route of infection.

Since the *Drosophila* innate immune system responds differently to Gram negative and Gram positive bacteria [61], we continued our pathogen challenges by microinjecting flies with *Staphylococcus aureus*. When flies were microinjected with ~100 CFUs, galbut virus infected DGRP 399 male flies survived slightly longer than their galbut virus infected counterparts (p=.021) as did DGRP 517 females (p=6.8×10^-4^). As for DCV challenge, although these effects were statistically significant, they were small in magnitude.

As a final pathogen challenge, we injected flies with ~500 cells of the fungal pathogen *Candida albicans* [43]. Both male and female DGRP 399 galbut virus infected flies died faster than their uninfected counterparts following *C. albicans* challenge (**Fig 6E**; DGRP female p=6.5×10^-6^ and DGRP male p=3.5×10^-5^). No significant differences were observed for DGRP 517 flies (**Fig 6E**).

### Galbut virus induces strain and sex specific changes in the transcriptome

We used RNA sequencing (RNA-seq) to explore transcriptional changes that could underlie the observed phenotypic differences between galbut virus infected and uninfected flies. We sequenced mRNA from pools of 10 whole adult females or males. We first performed hierarchical clustering to assess similarity between gene expression profiles in all datasets (**Fig. 7**). Sex was by far the most important variable influencing gene expression patterns: datasets from males and females were completely separated, with long branch lengths separating the clusters. This separation likely reflects the different chromosome repertoires of males and females in addition to sex-specific expression differences. Male datasets then clustered by DGRP strain, with DGRP 399 and 517 males forming separate subclusters. In DGRP 399 males, the group with the highest galbut virus RNA levels (**Fig. 1**), galbut virus infected and uninfected flies formed discrete clusters. In DGRP 517 males, the separation of infected and uninfected flies was not as clean. Gene expression patterns of female flies did not form subclusters based on DGRP strain nor galbut virus infection status (**Fig. 7**). Thus, as with other phenotypes, fly sex and DGRP strain influenced gene expression more than galbut virus infection.

**Fig 7.**
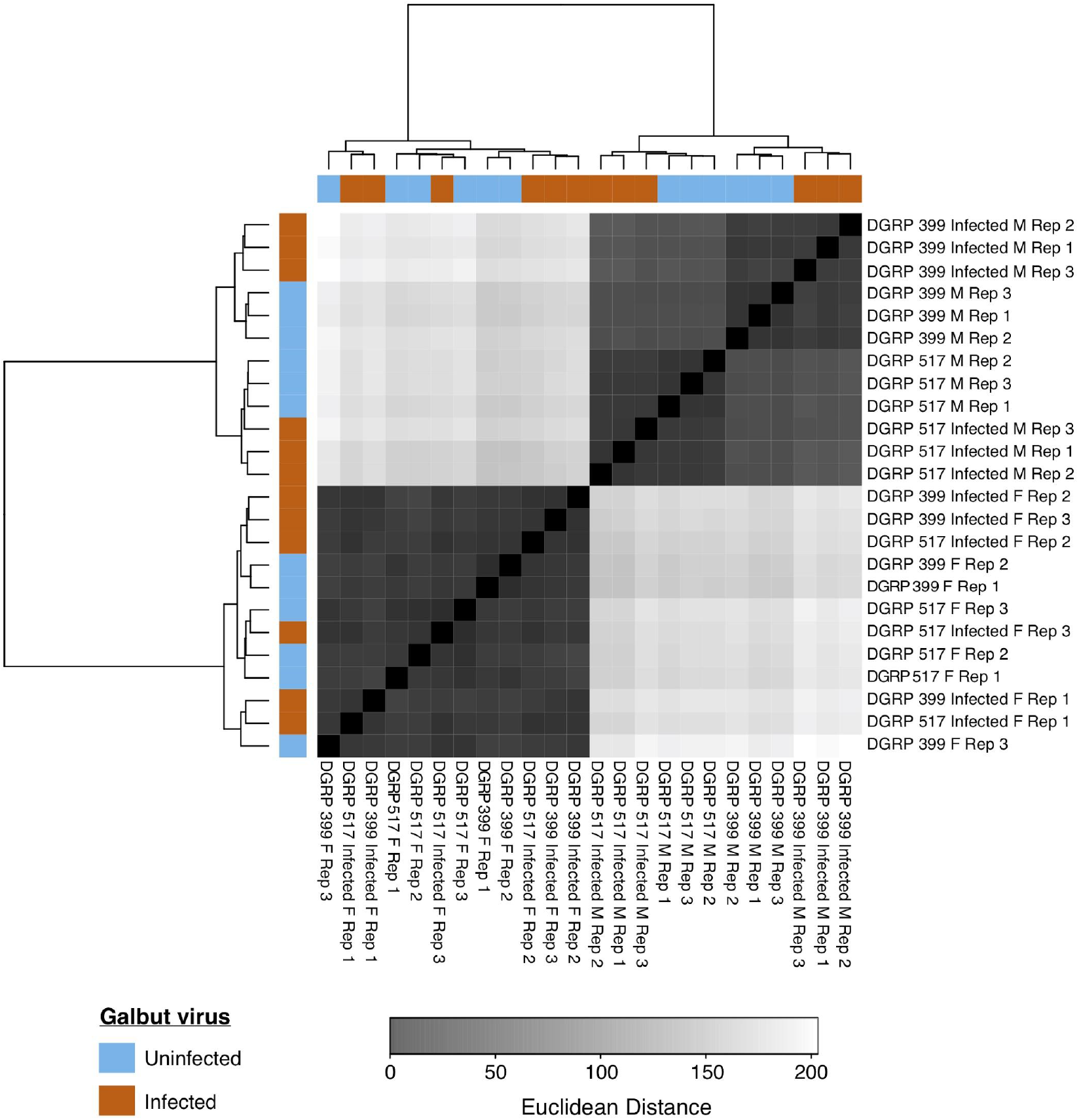
Galbut virus exerts minimal impact on overall transcriptional responses in flies. A sample distance matrix (Euclidean distances) quantifying the similarity between gene expression patterns in all datasets. Rep: biological replicates of 10 flies per replicate.

Only a single gene exhibited significant differential expression (adjusted p-value (padj) < 0.05) in all flies when compared by galbut virus infection status alone. The gene was a ribosomal RNA pseudogene (28S ribosomal RNA pseudogene CR45851) and it was upregulated in all groups of infected flies. We therefore examined transcriptional responses in flies grouped by DGRP strain and sex. Within these subsets, the response to galbut virus infection varied by both the number of differentially expressed genes and those that passed a significance threshold (**Fig 8**). Given a lack of consistent fitness phenotypes across any one sex or strain in most cases, this may be unsurprising.

**Fig 8.**
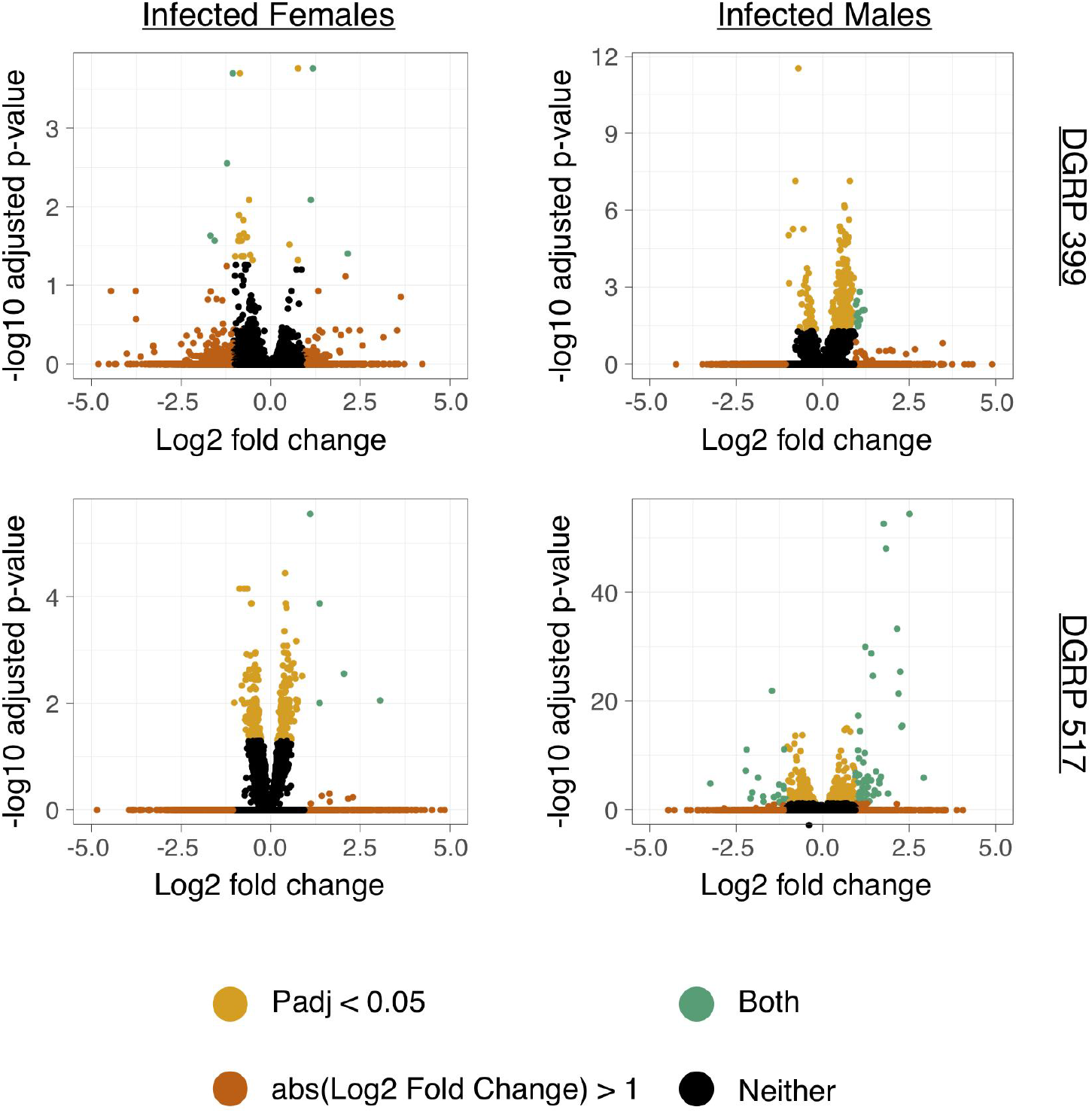
Volcano plots of differential gene expression in galbut virus infected flies. Plots depict the relative fold change of individual genes in galbut virus infected flies relative to uninfected flies (positive fold-change values indicate higher expression levels in galbut virus infected flies) on x axes and multiple testing corrected p-values on y axes. Individual genes that have a log2 fold change greater than 1 (orange), an adjusted p-value < 0.05 (gold), or both (green) are colored.

Among the top upregulated and downregulated genes in the experimental groups, few genes were shared between groups. Genes that were differentially expressed in more than one group included Kruppel homolog 1 (*Kr-h1*), which was significantly downregulated in galbut virus infected DGRP 399 and 517 females (**Fig 9**). This gene is a transcriptional regulator that has links to development [62–64]. Formin homology 2 domain containing (*Fhos*), which functions in development (remodeling of muscle cytoskeleton) and immune response (directs macrophage movement), was downregulated in both DGRP 399 infected females and males [65–67]. It is possible that these changes are related to the differences in developmental speed and pathogen susceptibility that we had observed (**Fig 4B; Fig 6B, E**).

**Fig 9.**
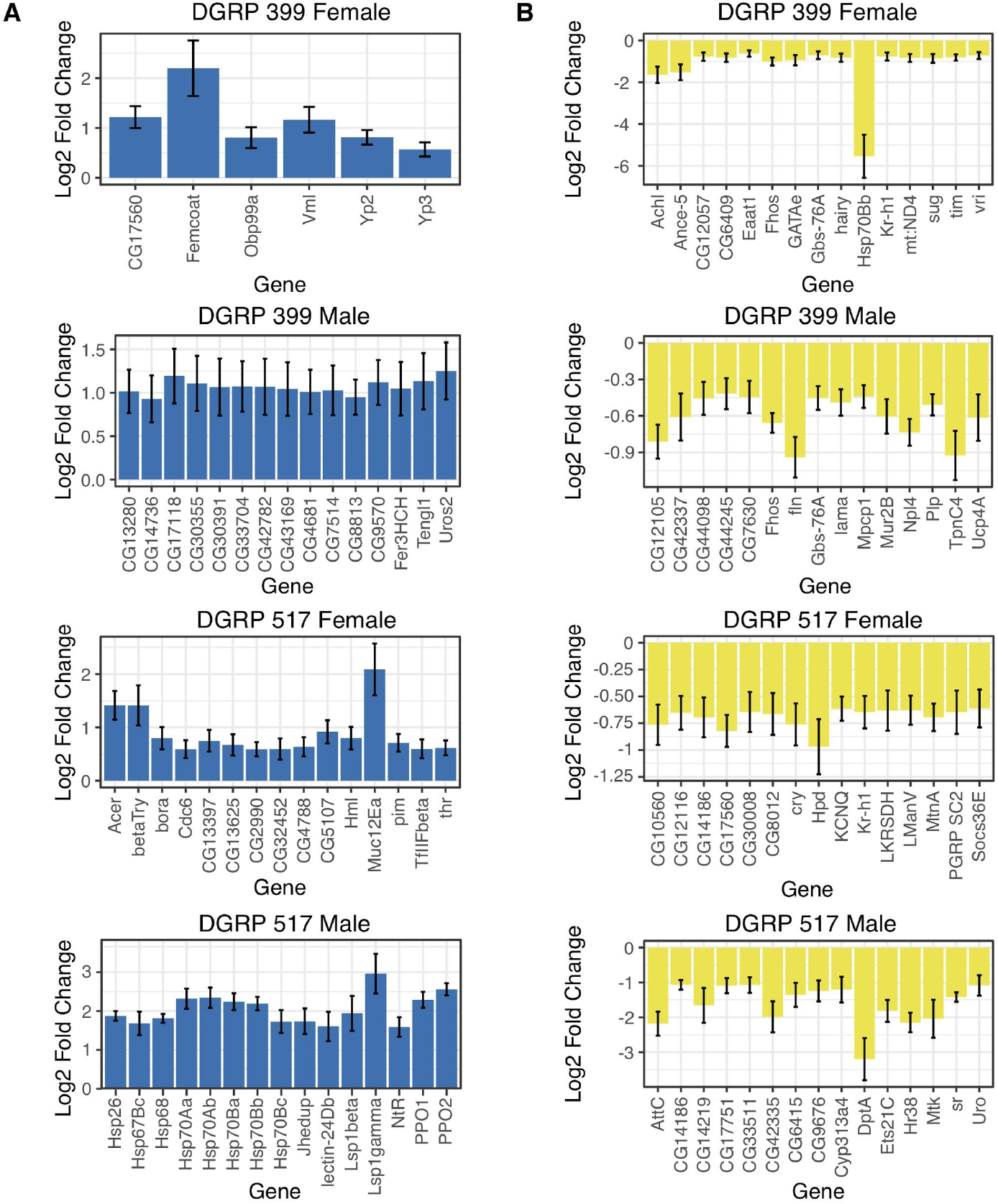
Top differentially expressed genes in infected flies relative to uninfected flies as. (A) Top 15 most significantly upregulated genes (ranked by padj, with padj < 0.05) in each experimental group relative to uninfected flies (B) Top 15 most significantly downregulated genes in each experimental group relative to uninfected flies.

Two genes with limited functional information were differentially expressed in both sexes of one or the other of the DGRP strains. Glycogen binding subunit 76A (*Gbs-76A*) was downregulated in galbut virus infected DGRP 399 flies of both sexes. This gene is inferred to play a role in the glycogen biosynthesis pathway [68]. In male and female DGRP 517 galbut virus infected flies, gene CG14186 was downregulated. CG14186 is affiliated with the biological process of cilium assembly, but its molecular function is unknown.

Two genes that were among the list of top differentially regulated genes across groups, but in opposite directions, were CG17560 and Heat shock protein 70Bb (*Hsp70Bb*). CG17560 is predicted to have implications in metabolic processes [68]. In DGRP 399 infected females, this gene was upregulated, while in DGRP 517 infected females, it was downregulated (**Fig 9**). *Hsp70Bb* was downregulated in DGRP 399 infected females, but was upregulated in DGRP 517 infected males (**Fig 9**). Heat shock proteins accounted for a large fraction of the upregulated genes in DGRP 517 infected males. Heat shock proteins are upregulated under heat and chemical stress, but these proteins have additional antiviral functions [69].

We performed a gene set enrichment analysis (GSEA) with the genes pre-ranked by log2 fold change using the clusterProfiler package in R [49] (**Figs S2–4, Tables 1–2**). Among the most significantly enriched gene ontology (GO) pathways (biological process ontologies only), pathways associated with development, morphogenesis, and metabolism were positively enriched in infected DGRP 517 flies (**Table 1, S2–3 Fig)**. GO pathways associated with neuron development and differentiation and response to stimuli were also differentially regulated (**Table 1, Figs S2–4**). GO pathways under the parent GO terms reproduction (GO:0000003) and reproductive process (GO:0022414) were positively enriched in galbut virus infected flies (**Tables 1 and 2**). Between 9 and 35 GO pathways associated with reproduction were enriched in galbut virus infected DGRP flies. All differentially regulated pathways, except 2 (DGRP 399 females, GO:0046008; DGRP 517 males, GO:0051446), were positively enriched in infected flies compared to uninfected flies.

**Table 1.**
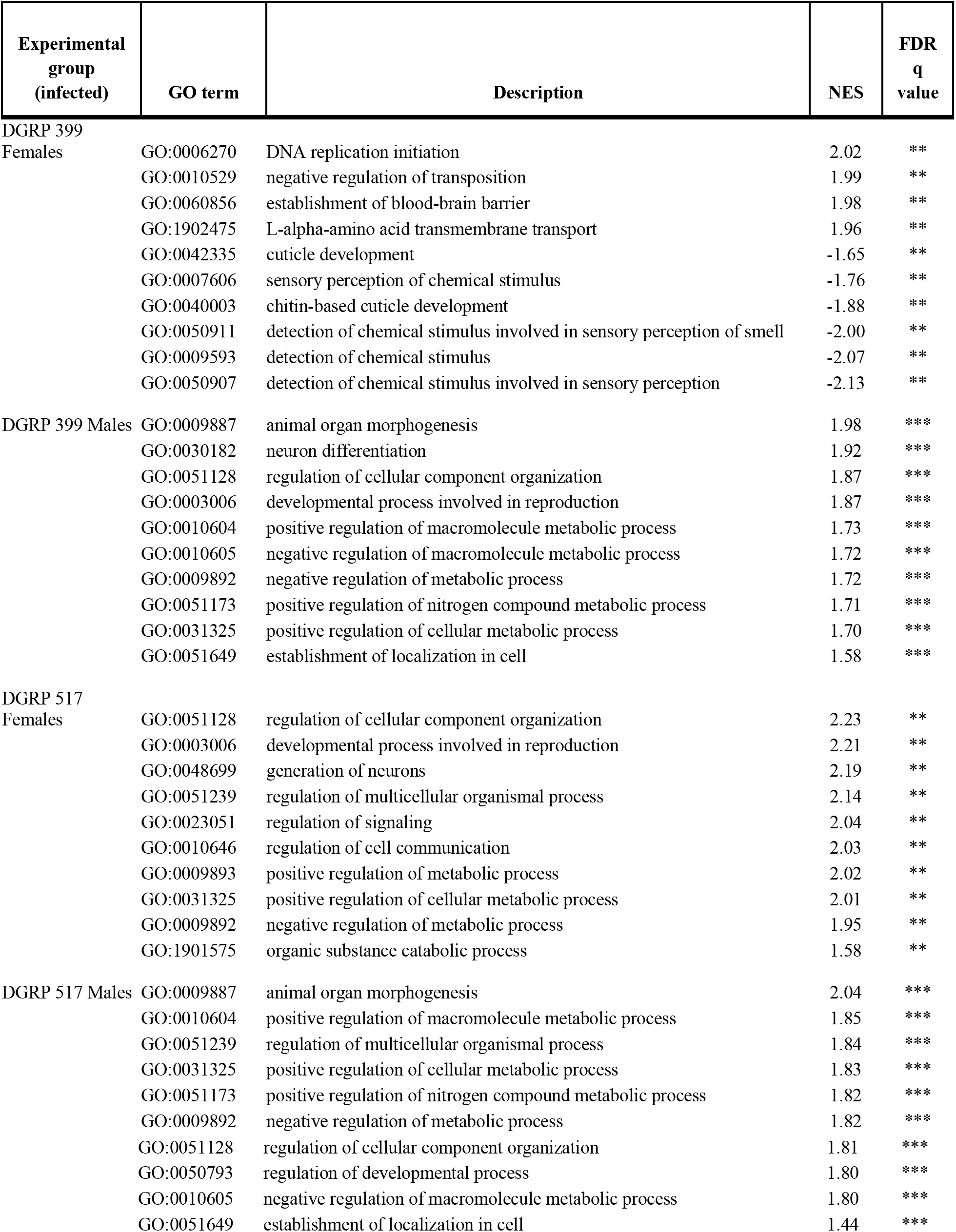
Most significantly enriched gene ontology pathways (biological process ontologies only) within infected flies identified via gene set enrichment analysis. NES: Normalized enrichment score; FDR: False Discovery Rate q value (Benjamini and Hochberg adjustment); *: q < 0.05, **: q < 0.01, ***: q < 0.001.

**Table 2.**
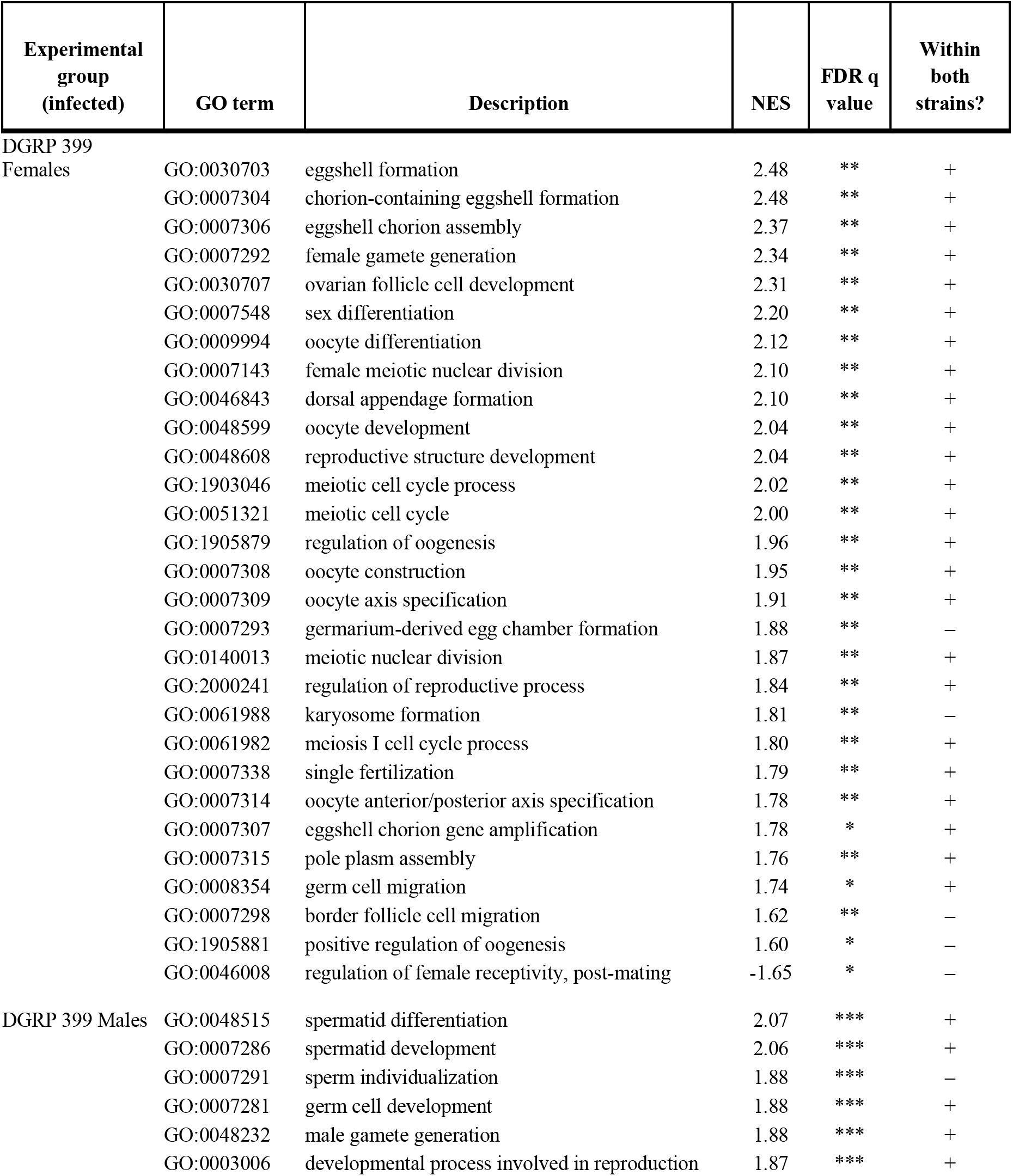
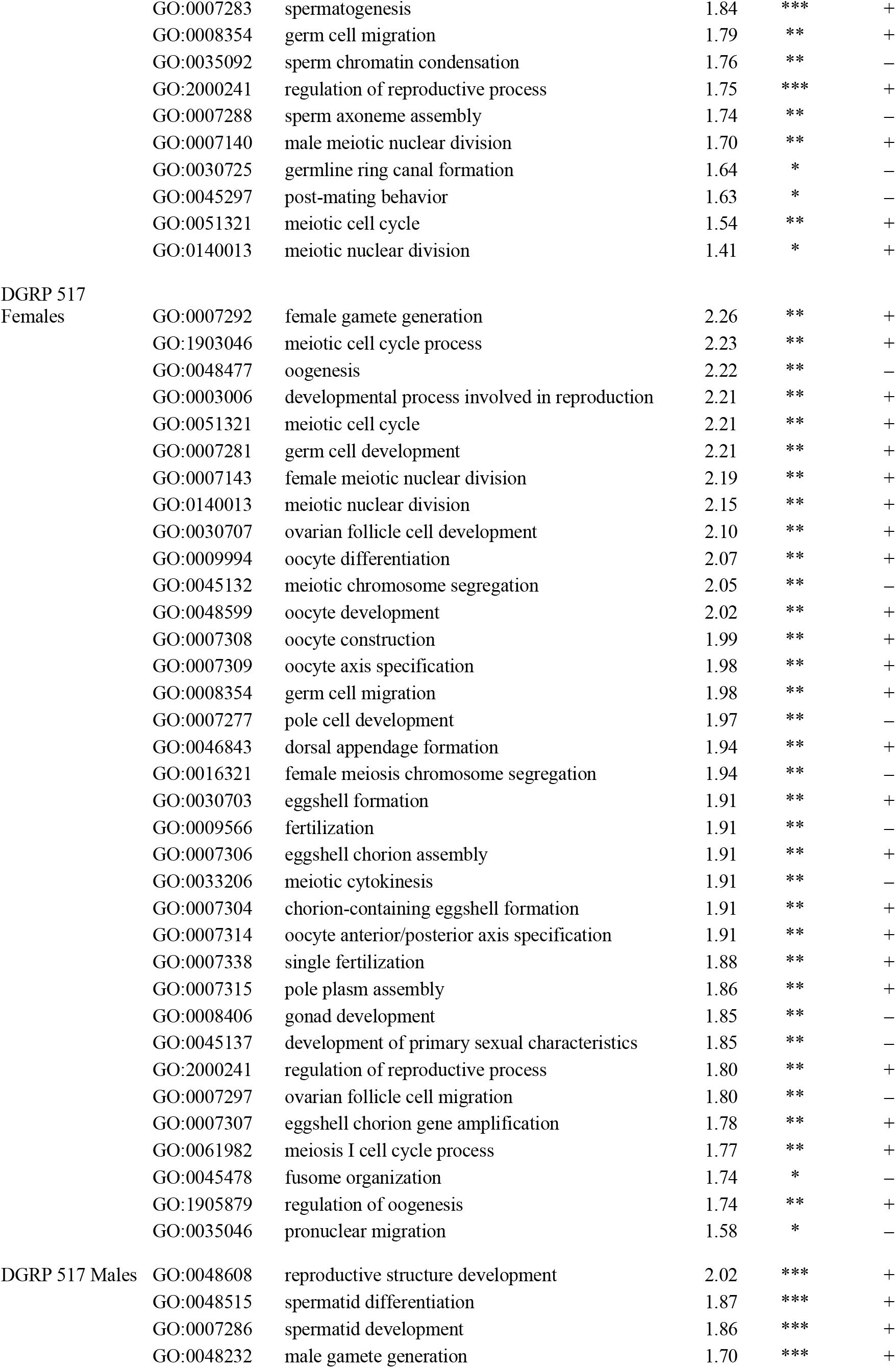
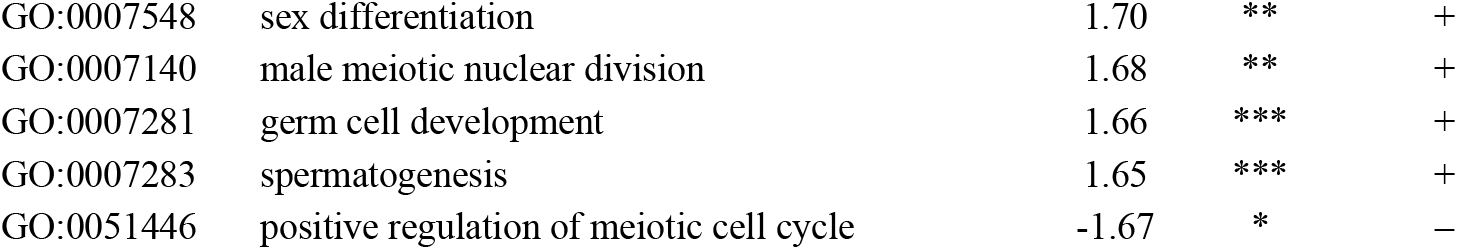
Enriched gene ontology pathways (biological process ontologies only) associated with reproduction identified via gene set enrichment analysis. NES: Normalized enrichment score; FDR: False Discovery Rate q value (Benjamini and Hochberg); *: q < 0.05, **: q < 0.01, ***: q < 0.001.

## Discussion

A major goal of this study was to understand why galbut virus, despite a high rate of vertical transmission (~100% from both parents), is maintained at a worldwide prevalence of only ~60%. We hypothesized that although galbut virus infection does not produce obvious phenotypic changes, infection might inflict enough of a fitness cost that resistant flies would experience a survival benefit. This would be analogous to an allele with a small negative selection coefficient. To test this hypothesis, we quantified multiple components of fitness in two genetic backgrounds.

Overall, galbut virus infection produced minimal measurable phenotypic effects. In some cases these would be predicted to decrease fitness, such as shortened average lifespan (**Fig. 3**) or decreased survival following fungal infection (**Fig 7E**). In other cases, trait differences such as faster development might increase the relative fitness of galbut virus infected flies (**Fig 5**).

Galbut virus infection minimally decreased lifespan and total offspring output, but the observed trends varied by DGRP strain and mostly did not rise to a statistically significant level (**Figs 3–4**). Our experiments lasted longer than the natural lifespan of *D. melanogaster*, which is estimated to be a week or less in the wild [70]. The total offspring output of galbut virus infected flies and uninfected flies only began to diverge after the parents were > 20 days old, and there was no impact on the number of eggs laid by young females over three days (**Fig 4**). Other examples of partitiviruses altering the reproductive output of their hosts include a partitivirus enhancing fecundity in *Cryptosporidium* [71], a reduction of spores from a partitivirus-infected fungus [72], and partitiviruses infecting *Spodoptera* moths that produced a major decrease in hatchling numbers [72].

Gene ontologies associated with reproduction were positively enriched in galbut virus infected flies, regardless of sex or strain (**Table 2**). It is possible that galbut virus infection manipulates reproductive pathways in a manner that contributes to efficient vertical transmission [4]. The upregulation of genes associated with oogenesis was observed in flies infected with *Drosophila melanogaster* sigma virus, which also depends on vertical transmission [73].

DGRP 517 flies infected by galbut virus pupated and reached adulthood faster than uninfected flies (**Fig 5**). An initial assumption would be that a faster developmental time, in combination with the short life of flies in the wild [70], would confer a fitness benefit. However, flies selected for faster development exhibited fitness trade-offs such as reduced body weight and size, decreased resistance to starvation and desiccation, and an overall lower egg output [74]. This highlights the difficulty of extrapolating total fitness from singly-measured traits [75].

For the most part, galbut virus infected and uninfected flies survived similarly following infection by microbial pathogens (**Fig 6**). Galbut virus infected DGRP 399 females exhibited decreased survival following ingestion, but not injection, of *Pseudomonas aeruginosa*. This difference may not be surprising as the gut epithelial immune response has key differences compared to responses to systemic infection [76]. DGRP 399 flies of both sexes exhibited increased sensitivity to the fungal pathogen *Candida albicans*. In *Drosophila*, the common microbiome constituent *Lactobacillus planatarum* decreased mortality of a fungal pathogen (*Diaporthe* sp.) by mitigating fungal toxicity and altered fly behaviour to reduce infection risk [77]. No significant changes in the DNA levels of *L. planatarum* or other major microbiome constituents was observed in galbut virus infected flies (**Fig 2**).

It is difficult to assess the net impact of these separately measured traits. Laboratory assays imperfectly recapitulate natural environments and these experiments provide a limited window into the influence of galbut virus in the wild. For most measured traits, differences associated with galbut virus infection were smaller than those attributable to different DGRP strain and sex (**Figs 3–4, Fig 7**). Nevertheless, selection can act on small differences in relative fitness, and it is possible that in aggregate galbut virus infection reduces fitness. Galbut virus is highly prevalent, exhibits a broad tissue distribution, and exists as a lifelong infection, so small phenotypic changes should not necessarily be interpreted as insignificant ones. Additional laboratory and field-based studies that track galbut virus-*Drosophila* dynamics will shed further light on the extent to which this virus and similar persistent viruses shape the evolution of their hosts in cryptic but possibly important ways.

## Acknowledgements

We thank Marylee Kapuscinski and the Colorado State University Next Generation Sequencing Core Facility for assistance with sequencing and Dr. Susan Tsunoda for helpful discussion. We would like to acknowledge Drs. Raul Andino and Michel Tassetto for providing *Drosophila* C virus stocks and Dr. Brad Borlee for providing *Pseudomonas aeruginosa* and *Staphylococcus aureus* stocks.

## SUPPLEMENTAL MATERIALS

**Supplemental Fig 1.**
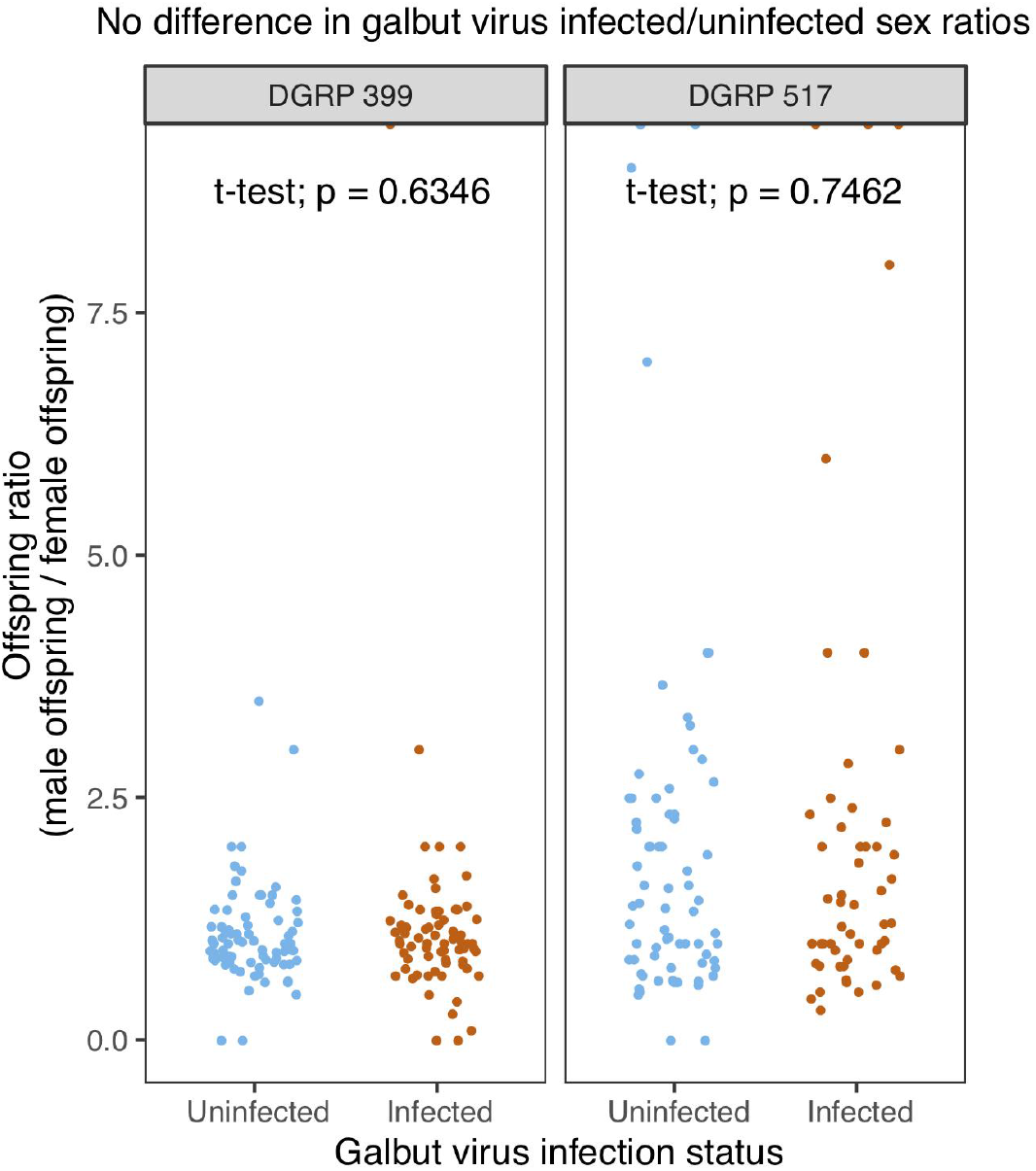
Galbut virus infection does not influence adult offspring sex ratio. Offspring collected from groups of galbut virus infected or uninfected parents from DGRP 399 and 517 strains every 14 days (see **Fig 4**). Offspring sex ratios from each time point were calculated by dividing total male offspring by total female offspring. No statistical significance was measured in either strain (t-test).

**Supplemental Fig 2.**
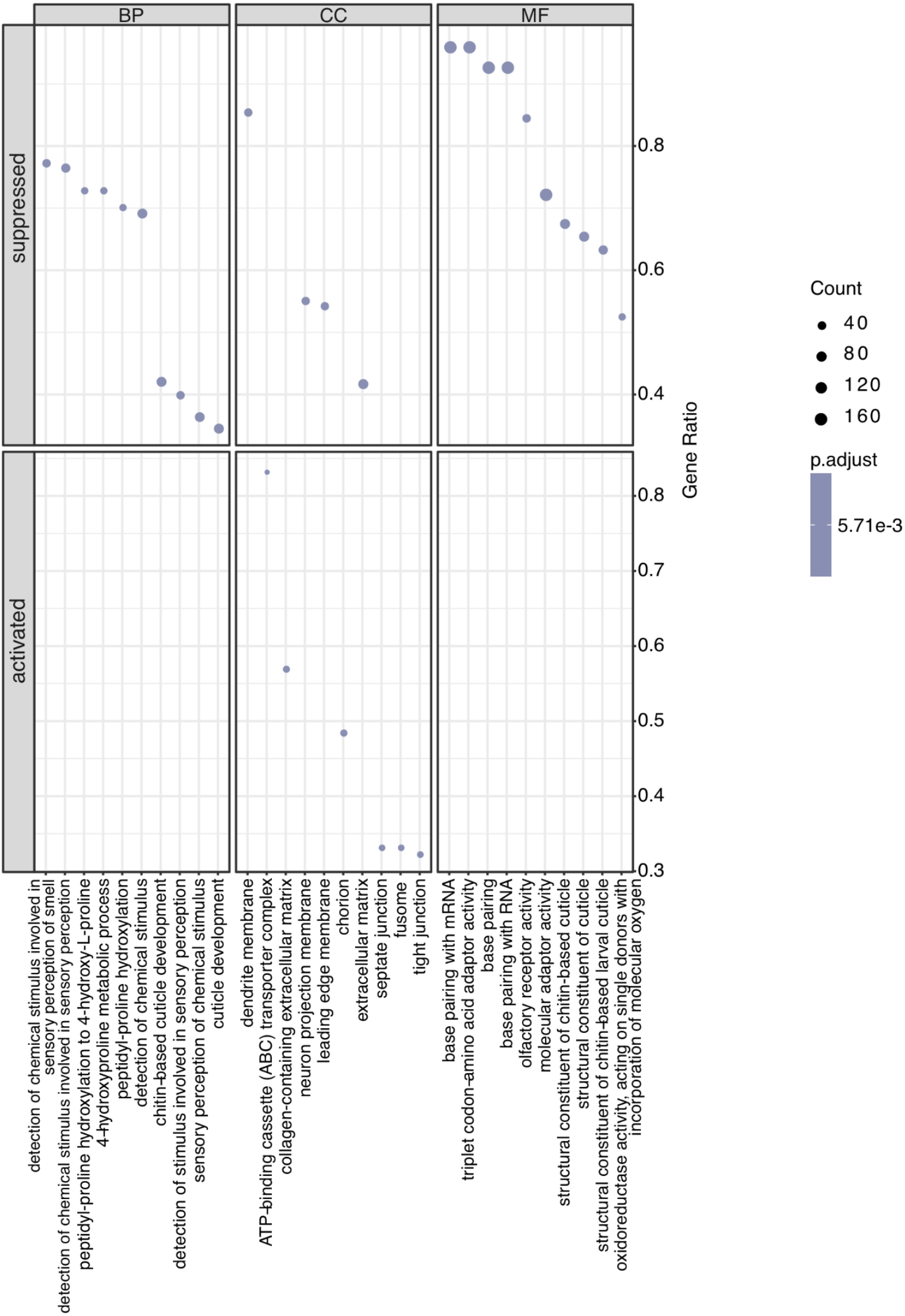
Dot plot of differentially regulated gene ontology (GO) pathways in infected DGRP 399 female flies. A dot plot representation of the top differentially regulated GO pathways in galbut virus-infected DGRP 399 female flies as determined by gene set enrichment analysis (GSEA) using the R package “clusterProfiler”. Top 10 differentially regulated pathways are plotted in each GO category (biological function, BF; cellular component, CC; molecular function, MF). Differentially regulated pathways for these flies were either upregulated (activated) or downregulated (suppressed). Size of dots corresponds with number of differentially regulated genes (DEG; counts) identified in each specified GO pathway. Percentage of DEGs in a given GO pathway (number of DEGs divided by total number of genes listed under the specified GO pathway) is plotted as gene ratio.

**Supplemental Fig 3.**
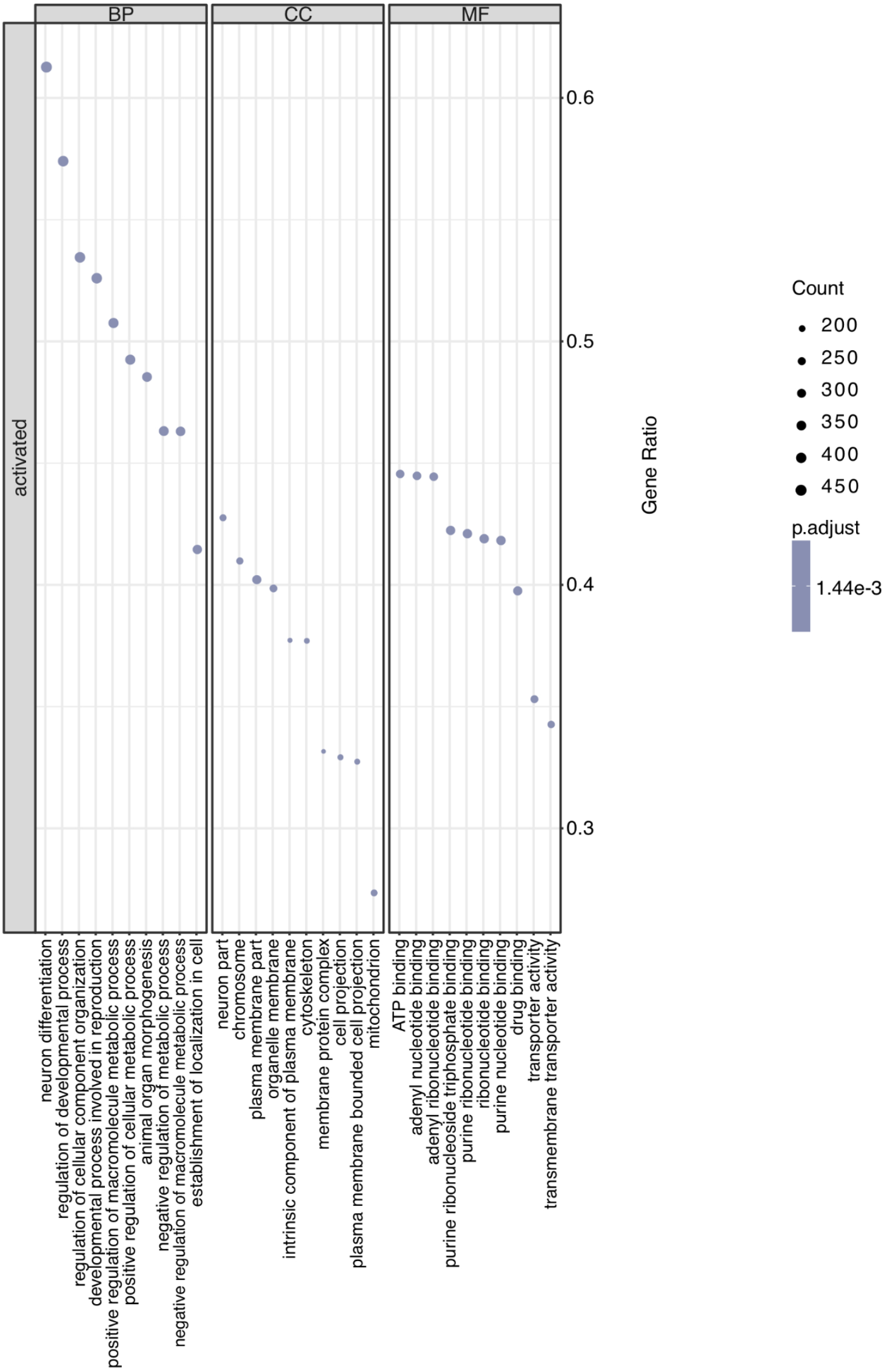
Dot plot of differentially regulated gene ontology (GO) pathways in infected DGRP 399 male flies. A dot plot representation of the top differentially regulated GO pathways in galbut virus-infected DGRP 399 male flies as determined by gene set enrichment analysis (GSEA) using the R package “clusterProfiler”. Top 10 differentially regulated pathways are plotted in each GO category (biological function, BF; cellular component, CC; molecular function, MF). All top differentially regulated pathways for these flies were upregulated (activated). Size of dots corresponds with number of differentially regulated genes (DEG; counts) identified in each specified GO pathway. Percentage of DEGs in a given GO pathway (number of DEGs divided by total number of genes listed the specified GO pathway) is plotted as gene ratio.

**Supplemental Fig 4.**
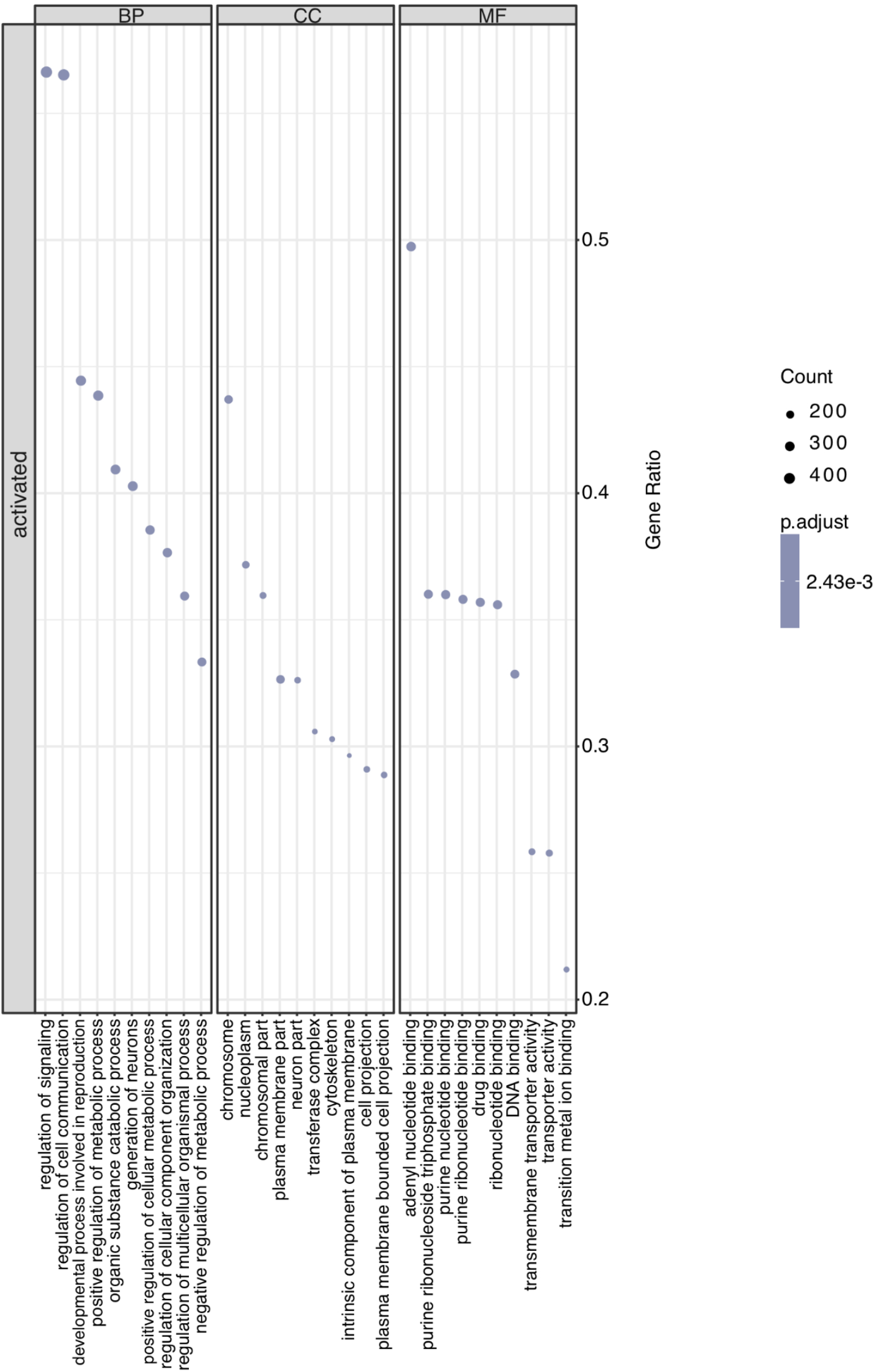
Dot plot of differentially regulated gene ontology (GO) pathways in infected DGRP 517 female flies. A dot plot representation of the top differentially regulated GO pathways in galbut virus infected DGRP 517 female flies as determined by gene set enrichment analysis (GSEA) using the R package “clusterProfiler”. Top 10 differentially regulated pathways are plotted in each GO category (biological function, BF; cellular component, CC; molecular function, MF). All top differentially regulated pathways for these flies were upregulated (activated). Size of dots corresponds with number of differentially regulated genes (DEG; counts) identified in each specified GO pathway. Percentage of DEGs in a given GO pathway (number of DEGs divided by total number of genes listed under the specified GO pathway) is plotted as gene ratio.

**Supplemental Fig 5.**
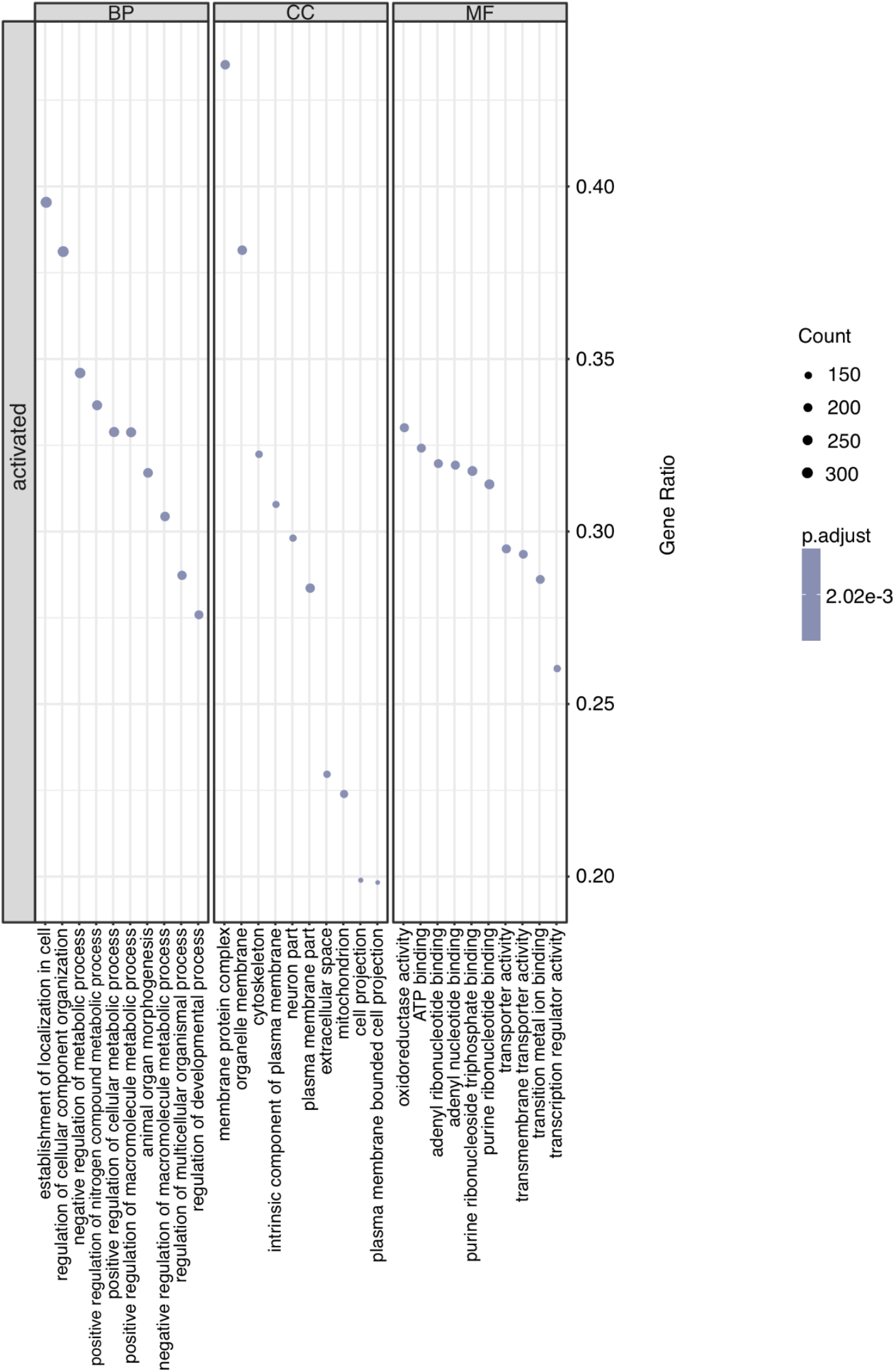
Dot plot of differentially regulated gene ontology (GO) pathways in infected DGRP 517 male flies. A dot plot representation of the top differentially regulated GO pathways in galbut virus-infected DGRP 517 male flies as determined by gene set enrichment analysis (GSEA) using the R package “clusterProfiler”. Top 10 differentially regulated pathways are plotted in each GO category (biological function, BF; cellular component, CC; molecular function, MF). All top differentially regulated pathways for these flies were upregulated (activated). Size of dots corresponds with number of differentially regulated genes (DEG; counts) identified in each specified GO pathway. Percentage of DEGs in a given GO pathway (number of DEGs divided by total number of genes listed under the specified GO pathway) is plotted as gene ratio.

**Supplemental Table 1:**
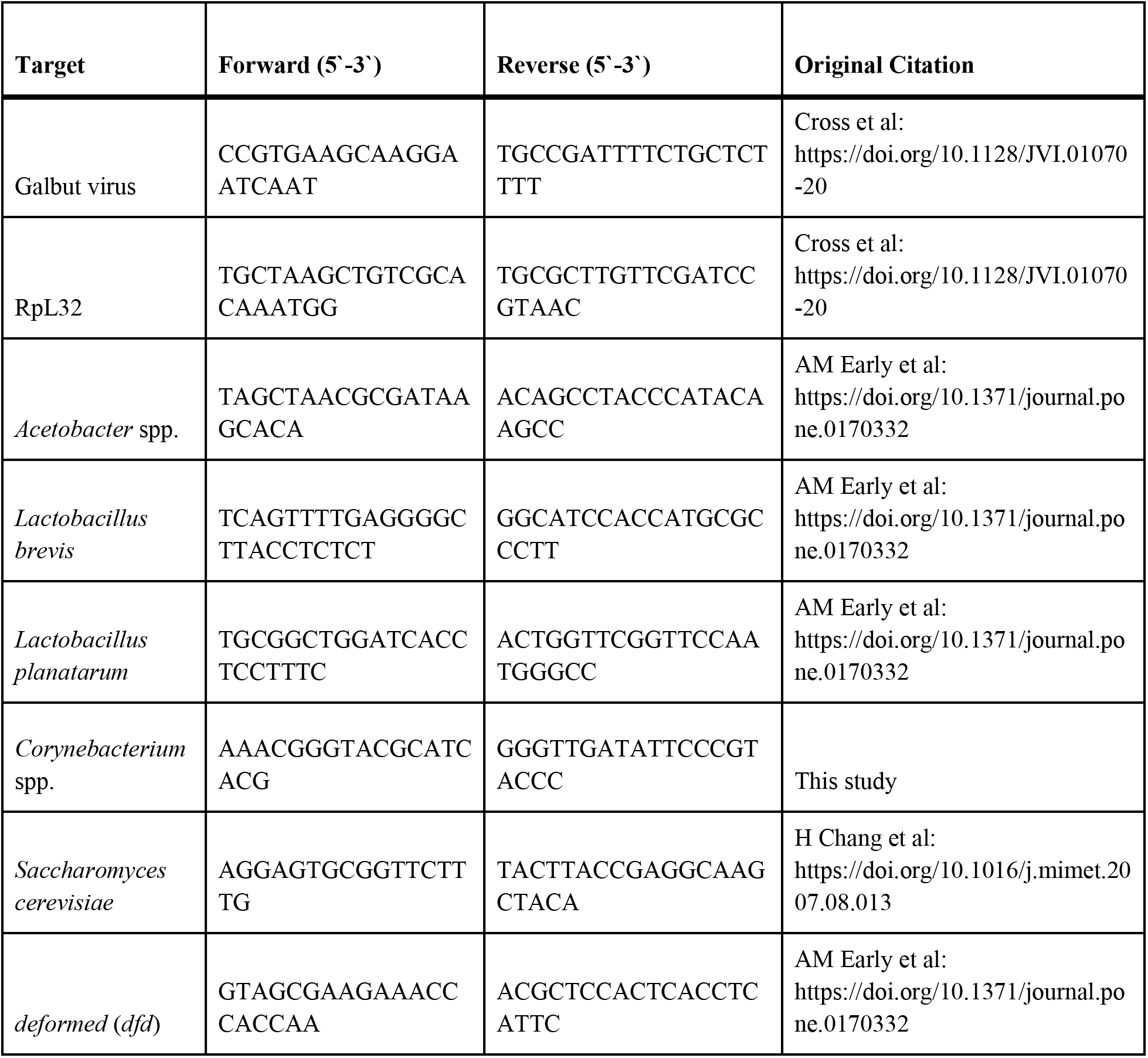
Primers used for quantifying levels of galbut virus and microbiome constituents.

